# A heterochromatin domain forms gradually at a new telomere and is highly dynamic at stable telomeres

**DOI:** 10.1101/211565

**Authors:** Jinyu Wang, Jessica R Eisenstatt, Julien Audry, Kristen Cornelius, Matthew Shaughnessy, Kathleen L Berkner, Kurt W Runge

**Affiliations:** Department of Genetics and Genome Sciences, Case Western Reserve University, Cleveland, OH, USA; Department of Biochemistry, Case Western Reserve University, Cleveland, OH, USA; Departments of Immunology and Molecular Genetics, Lerner Research Institute, Cleveland Clinic Foundation, Cleveland, OH, USA; Department of Molecular Genetics, The Ohio State University, Columbus, OH, USA

## Abstract

Heterochromatin domains play important roles in chromosome biology, organismal development and aging. In the fission yeast *Schizosaccharomyces pombe* and metazoans, heterochromatin is marked by histone H3 lysine 9 dimethylation. While factors required for heterochromatin have been identified, the dynamics of heterochromatin formation are poorly understood. Telomeres convert adjacent chromatin into heterochromatin. To form a new heterochromatic region in *S. pombe*, an inducible DNA double-strand break (DSB) was engineered next to 48 bp of telomere repeats in euchromatin, which caused formation of new telomere and gradual spreading of heterochromatin. However, spreading was highly dynamic even after the telomere had reached its stable length. The system also revealed the presence of repeats located at the boundaries of euchromatin and heterochromatin that are oriented to allow the efficient healing of a euchromatic DSB to cap the chromosome end with a new telomere. Telomere formation in *S. pombe* therefore reveals novel aspects of heterochromatin dynamics and the presence of failsafe mechanisms to repair subtelomeric breaks, with implications for similar processes in metazoan genomes.

## Introduction

A central question in eukaryotic biology is the establishment and maintenance of chromatin domains, i.e. regions of nucleosomal DNA where the histone composition and spectrum of post-translational modifications are similar. As embryonic cells differentiate, cell type-specific gene expression is established in part by the establishment and maintenance of chromatin domains (e.g. changes in the globin locus in hematopoietic cells, X-chromosome inactivation in females mammals (1, 2)). Chromatin domain reorganization also occurs during tumorigenesis as cells transform into rapidly growing cancers (3). Heterochromatin domains, marked in part by nucleosomes with di- and tri-methylation of lysine 9 of histone H3 (H3K9me2 or 3), have been intensely studied for their role in chromosome biology. Heterochromatin domains are known for silencing gene expression (4), and can be induced during mammalian cell senescence and aging to form senescence-associated heterochromatin foci containing H3K9me2 (5–7). Heterochromatin also plays an important role at centromeres, the chromosomal structure required for chromosome segregation at mitosis, as centromeric chromatin is flanked by heterochromatin domains that are required for complete function (8–11). While many of the factors required to maintain heterochromatin have been identified, the dynamics of how heterochromatin domains assemble and disassemble have remained long-standing, major questions that are only now being investigated (12–14).

Telomeres, the physical ends of chromosomes, are a second chromosomal structure bordered by heterochromatin. In yeast, humans and many other eukaryotes, telomeres consist of simple DNA repeats bound by specific proteins. These repeats and their associated proteins provide the first discovered essential function of telomeres: “that of sealing the end of the chromosome” (15), and distinguishing it from a Double-Strand Break (DSB)(15, 16). The second essential function is to replace sequences lost due to incomplete replication, which is accomplished by repeat addition via telomerase ((17, 18), reviewed in (19)). Telomeres also alter the adjacent nucleosomal chromatin to silence the expression of nearby genes (20, 21). However, as mutations that eliminate silencing do not cause telomeres to behave as DSBs (20, 22, 23), the essential functions of telomeres act independently of gene silencing. Heterochromatic gene silencing is associated with the presence of H3K9me2 in humans, flies and the fission yeast *Schizosaccharomyces pombe* (24), and *S. pombe* telomere-associated chromatin has the H3K9me2 modification (22, 23, 25). Thus, *S. pombe* telomeres provide an ideal model system to study heterochromatin and heterochromatin dynamics.

A major difficulty that impedes the investigation of heterochromatin domain dynamics is the large amount of time between the initiation of domain formation and its analysis. Telomeres have been formed in *S. pombe* by integrating *in vitro* constructed telomeric DNA into genomic sites *in vivo*, but 30 population doublings (PDs) or more must pass between the formation of the new telomere and the production of enough cells for chromatin and phenotypic analysis (21). Similar approaches requiring many PDs have followed heterochromatin formation at centromeres and other loci by introducing wild type genes into mutants defective for heterochromatin assembly (26–29). This approach also converts a mutant cell to a wild type one, so the levels of cellular chromatin proteins during domain formation are initially different than wild type cells. Consequently, the kinetics of heterochromatin formation and how the H3K9me2 modification spreads from the initiating site into the surrounding chromatin is largely unknown in either wild type or mutant cells. One hypothesis would be that spreading occurs immediately after the initiating site is created and quickly establishes the final heterochromatin domain within one or two generations (as with Sir protein spreading in *Saccharomyces cerevisiae* (30–32)). Alternatively, the formation of the initiation site, e.g. a functional telomere, may allow spreading over many cell divisions, with the size of the heterochromatin domain gradually increasing over time to form the final state (as suggested for several histone modifications in (30) and in *S. pombe* in (14)).

An inducible telomere formation system would provide an approach to study the kinetics of heterochromatin formation in wild type cells. Such systems contain a selectable marker followed by an internal tract of telomere repeats and a unique restriction site or cut site not present elsewhere in the genome. By placing the restriction enzyme or endonuclease gene under the control of a rapidly inducible promoter, one can induce a DSB in a large population of cells to expose the telomere repeats at the new chromosome end (33). In *S. cerevisiae* and *S. pombe*, a DSB in the middle of a chromosome normally leads to DNA degradation and growth inhibition (33–35)(Figure 1A). In contrast, a telomere formation system in *S. cerevisiae* and mammalian cells has shown that a DSB which exposes telomere repeats is immediately converted into a short, functional telomere that is not degraded (33, 36–38)(Figure 1B). The *S. cerevisiae* telomere formation system has yielded important insights into the roles of telomerase, telomere binding proteins, DNA polymerases and DNA damage proteins in telomere elongation (reviewed in (39)). However, *S. cerevisiae* lacks the H3K9me2 modification system, and so its use in modeling the kinetics of heterochromatin spreading that occurs in metazoans is limited.

**Figure 1.**
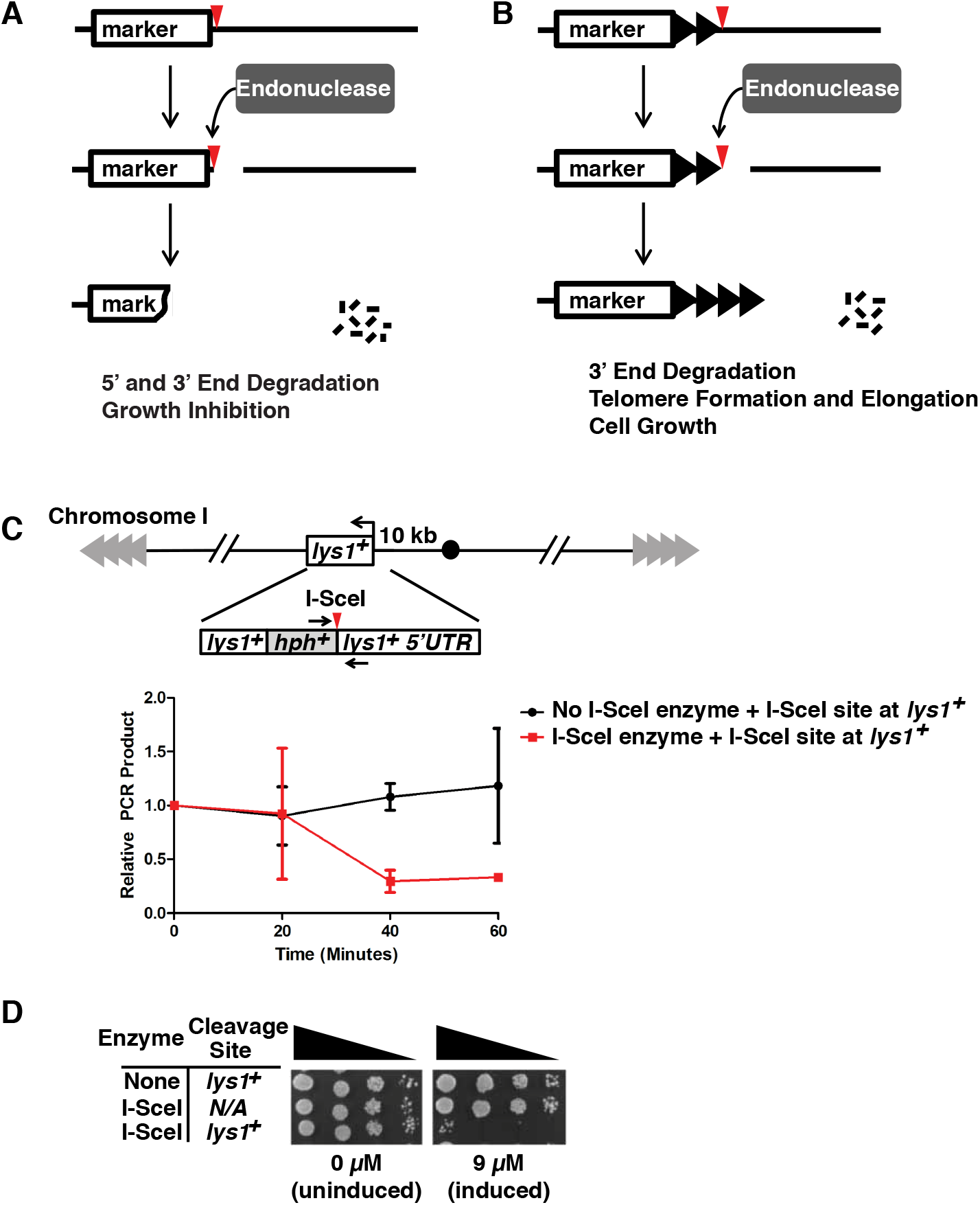
Double-strand break (DSB) systems and rapid I-SceI DSB formation. **(A) Inducible DSB system.** A restriction enzyme/endonuclease with no natural sites in the genome is produced in cells from a rapidly inducible promoter. After addition of the inducer and production of the endonuclease, a single site introduced into the genome (red triangle) can be cut to produce a DSB. In the DSB system, the strands both 5’ and 3’ to the endonuclease site are degraded (indicated by small black lines and loss of the marker DNA) and cell growth is inhibited. **(B) Inducible telomere formation system.** The new DSB exposes telomere repeats (black triangles) to form a new functional telomere that is stable and elongated. If the chromosomal sequences 3’ to the endonuclease site are dispensable, the new functional telomere allows normal cell growth. **(C) Rapid induction of an I-SceI-generated DSB.** The I-SceI restriction site was inserted into the 5’ UTR of *lys1*^*+*^, a gene located 10 kb from the centromere of chromosome I. The expression of I-SceI was induced by the addition of ahTET to 9 µM. Cell samples were taken before (0 min) and after ahTET addition in 20 min intervals. Genomic DNA was prepared and assayed for cleavage at the I-SceI site by qPCR using primers across the site (denoted by black arrows) and normalizing to the production of a similarly sized fragment at *his3*^+^ (as in (34)). The average values of two independent experiments (where each qPCR is performed in triplicate for each experiment) and the SEM are shown. About 75% of the sites are cleaved by 40 min in this assay. We note that this assay cannot distinguish between sites that were never cut and those that were cut and then ligated back together with or without mutation of the site. **(D) The I-SceI DSB causes growth arrest.** Five-fold serial dilutions of cells bearing either the *lys1*^*+*^ allele with or without the I-SceI site or the expression vector with or without the I-SceI gene were spotted onto rich medium with either 0 or 9 µM ahTET. Only those cells with both the I-SceI expression vector and the I-SceI site have the capability of producing a DSB, and these cells showed the growth inhibition associated with DSB induction.

*S. pombe* is a useful model for studying the H3K9me2 heterochromatin system (40), but a telomere formation system was previously not feasible owing to the lack of a method to rapidly induce a DSB. Two different rapidly inducible systems have recently been established by Watson *et al.* using the HO endonuclease (41) and by ourselves using I-PpoI endonuclease (34). Unfortunately, neither system was well suited for inducing telomere formation. The HO system uses an *urg1*^+^ promoter that is induced by the addition of uracil, which interferes with the use of the *ura4*^+^ selectable marker. Expression of the *ura4*^+^ gene can be selected for or against, which allows the facile monitoring of expression by cell growth and has been a mainstay of gene silencing studies (21, 42, 43). Our I-PpoI system avoids this *urg1^+^* limitation by using an anhydrotetracycline (ahTET)-inducible promoter, but I-PpoI cuts in the rDNA of almost all eukaryotes, so strains bearing mutated rDNA repeats must be used. We therefore designed a new method to rapidly induce a single DSB in the *S. pombe* genome and used it to create a telomere formation system. Telomere formation was induced in a population of cells to follow heterochromatin formation in real-time. While a functional telomere formed immediately, the H3K9me2 modification spread slowly from the functional telomere over several generations. Surprisingly, the extent of spreading varied with growth conditions and over time even when the length of the telomere repeat tracts was constant. Thus, the established heterochromatin domain was surprisingly dynamic. We also discovered that a DSB in the euchromatin that lacks telomere repeats was rapidly healed with high efficiency when present near a telomere, in contrast to breaks in the middle of the chromosome. Therefore, the structure of the *S. pombe* genome contains an unanticipated failsafe mechanism to rescue telomere loss. These results in *S. pombe* suggest similar novel processes may also occur at metazoan telomeres and heterochromatin domains.

## Results

### The *S. pombe* telomere formation system

We first developed an inducible DSB system in *S. pombe* using the I-SceI homing endonuclease. I-SceI has no endogenous sites in the *S. pombe* genome (44) and the I-SceI system has the advantage over other DSB inducing systems by leaving the *ura4*^+^ selection, a common telomere silencing marker (21), intact and not requiring special strain backgrounds (34, 41). I-SceI has the disadvantage of inefficient and slow cutting (45, 46). We therefore designed an I-SceI gene with preferred *S. pombe* codons (47) and two nuclear localization signals (NLS) at the N-terminus to enhance expression and genomic DNA cleavage. This I-SceI variant was expressed from a TetR repressed promoter, which allows expression of the desired gene after addition of ahTET (Figure 2A). Cutting efficiency was tested in a strain with the I-SceI site at a marker gene near the centromere of chromosome I, *lys1*^+^ (Figure 1C). Most I-SceI sites were cut within 40 minutes of induction of I-SceI expression (Figure 1C). When plated on inducing medium, the strain expressing I-SceI and containing a site at *lys1*^+^ showed a severe growth defect (Figure 1D), as seen with other strains that continuously induce a DSB (34, 48–50).

**Figure 2.**
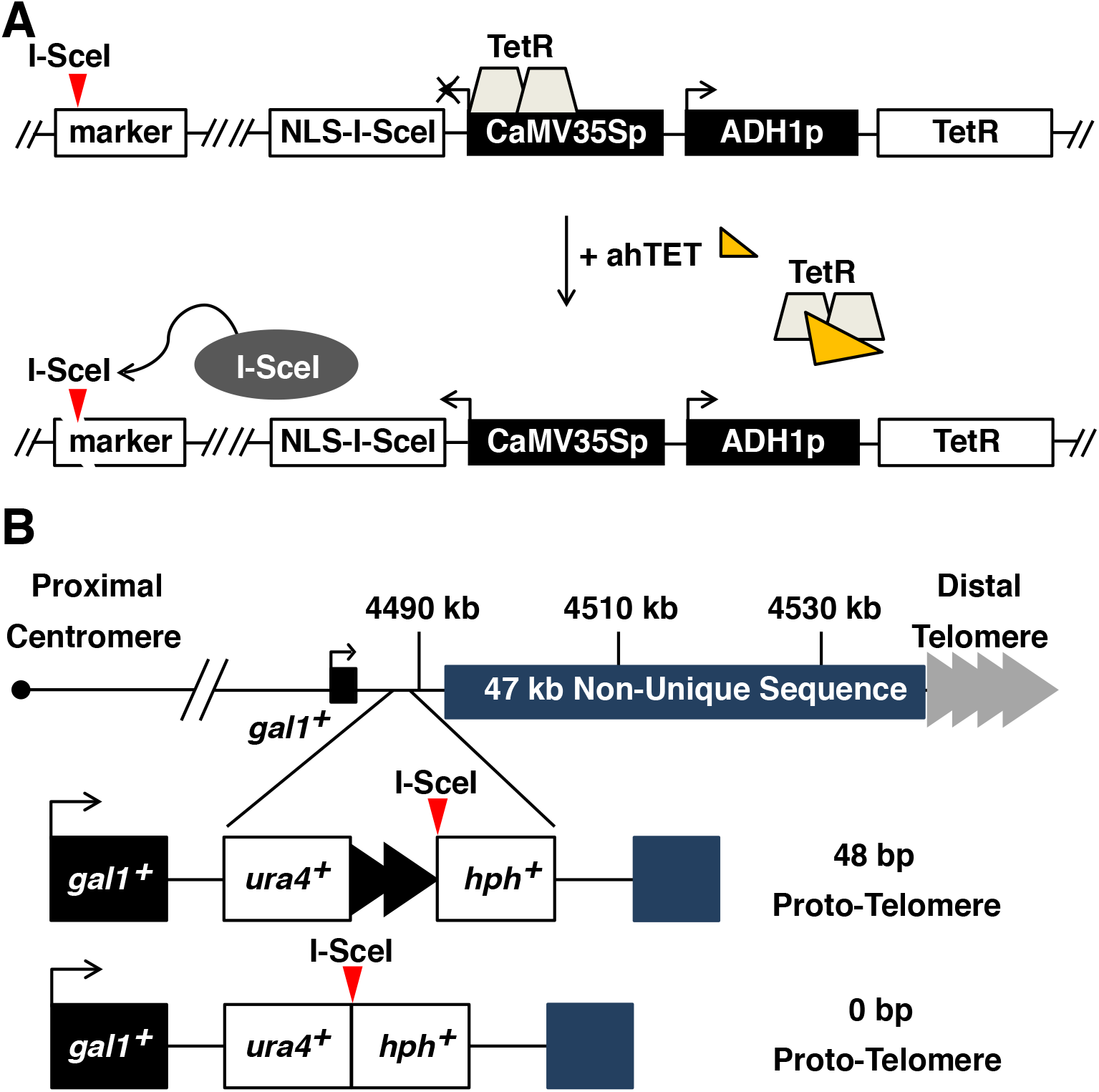
The I-SceI telomere formation system. **(A)** The I-SceI endonuclease was expressed from the tetracycline repressor (TetR) controlled CaMV35S promoter in a cassette that also expresses TetR. The addition of anhydrotetracycline (ahTET) induces I-SceI expression, which then cuts at sites introduced into the genome (red triangle). **(B)** The 48 bp proto-telomere contains the *ura4*^+^ gene followed by 48 bp of telomere repeats (black triangles) and the hygromycin resistance marker (*hph*^+^), while the 0 bp proto-telomere control lacks the telomere repeats. Both cassettes were inserted into the unique DNA 3’ of the *gal1*^+^ gene. The *S. pombe* endogenous telomere repeat tracts are indicated by grey triangles.

Two “proto-telomere” cassettes were created that contain either 48 bp or 0 bp of *S. pombe* telomere repeat sequence, an I-SceI site and two flanking selectable markers (Figure 2B). Cleavage at the I-SceI site should expose the telomere repeats and cause a loss of the distal marker and chromosome end. Consequently, to yield viable cells for analysis, the lost region had to be dispensable and the 48 bp telomere repeats must form a new functional telomere. We therefore chose a site in the 2 kb region 3’ to the *gal1*^+^ gene on the right end of chromosome II (IIR), because this region is unique in the genome and borders a 47 kb subtelomere-containing sequence that is repeated at both ends of chromosomes I and II (44, 51, 52). Cells that have lost most of these subtelomeric sequences are viable (52). In addition, *gal1*^+^ and each gene in the 86 kb region 5’ to *gal1*^+^ are not required for growth. Therefore, cleavage in the 2 kb region 3’ to *gal1*^+^ would cause loss of dispensable chromosomal sequences and allow the formation of a large heterochromatic domain near the proto-telomere after I-SceI cutting was induced and still produce viable cells.

### A functional short telomere forms after I-SceI cleavage

Telomere formation was induced in *S. pombe* cells containing the 48 bp proto-telomere by expressing I-SceI and monitoring the fate of the *ura4*^+^ and *hph*^+^ proto-telomere fragments by Southern analysis (Figure 3A). The uncut *ura4^+^-telomere repeats-hph^+^* band was visible prior to induction of I-SceI, and was replaced by the I-SceI cleaved *ura4*^+^ and *hph*^+^ bands over time. The *ura4*^+^ fragment was stable and increased in size and heterogeneity during the experiment, as expected for the elongation of the exposed 48 bp telomere repeats (Figure 3A). Elongation was almost certainly by telomerase as sequencing of the telomere fragments revealed that addition of new telomeric repeats occurred, in all but one case, to the I-SceI cleaved proto-telomere (Figure 4A), consistent with telomerase-mediated addition events in *S. cerevisiae* and mammalian cells (38, 53–56). When telomere formation was performed in cells lacking telomerase RNA, the newly formed telomere was stable but not elongated (Figure 5). In contrast, the *hph*^+^ fragment was rapidly degraded (Figure 3A). Thus, formation of a telomere, the stable structure that “seals” the end of the chromosome (15), occurs at the earliest time point tested and is independent of telomerase activity. Following elongation of the telomere repeats for over 50 PDs revealed that the telomere repeat tracts were stably maintained (Figure 4B) and reached their equilibrium final lengths by ~8 PDs (Figure 4C). The initial stability and subsequent elongation of the *ura4*^+^*-48 bp telomere* band therefore show that this fragment rapidly acquired the essential telomeric functions of end-capping and end-replication after I-SceI cleavage, and behaved the same as short functional telomeres at chromosome ends (e.g. newly formed *S. cerevisiae* telomeres and telomeres of cells lacking Tel1, MRX or Ku (20, 57)).

**Figure 3.**
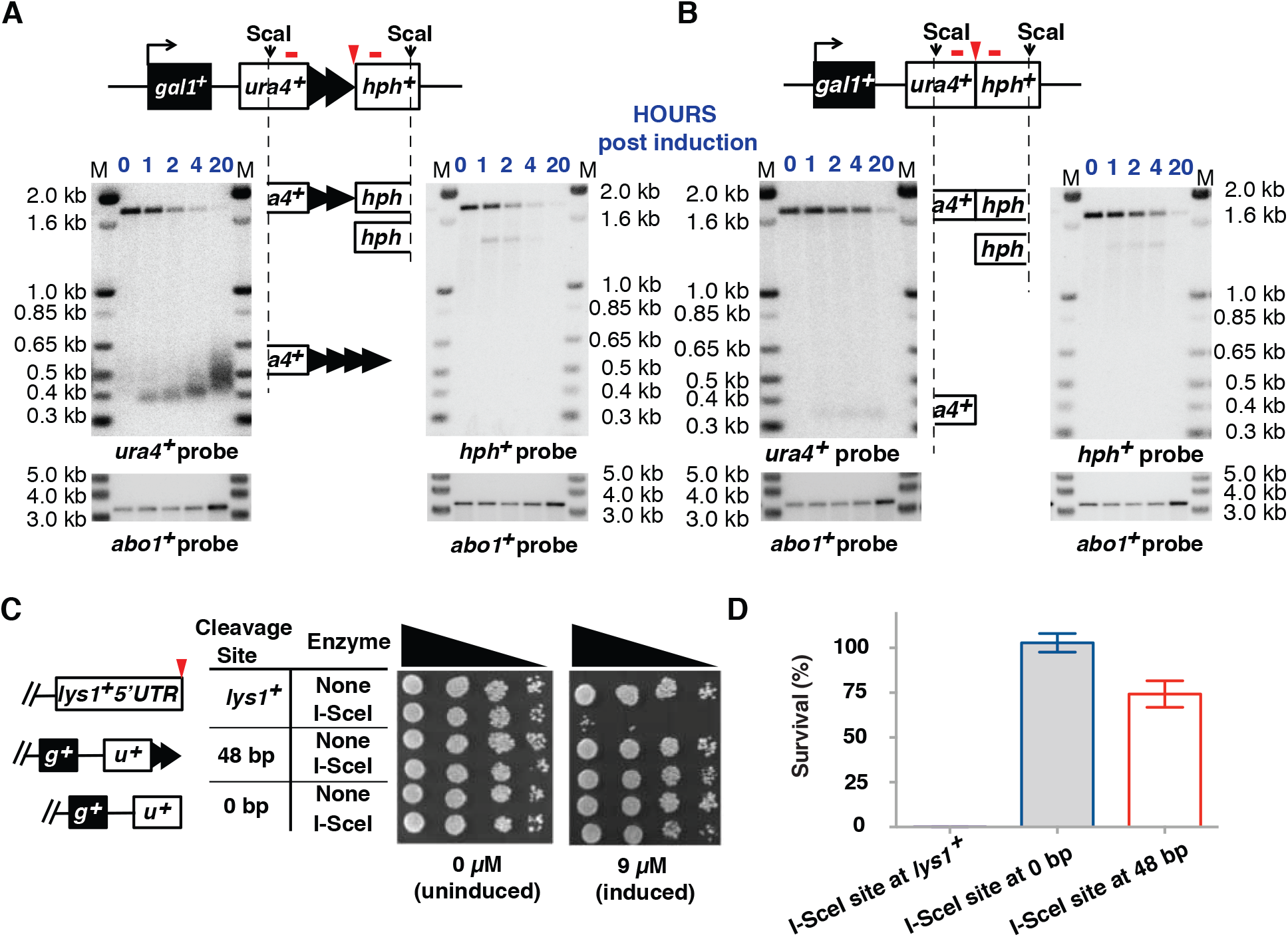
I-SceI cleavage converts the 48 bp proto-telomere to a telomere. **(A)** Exponentially growing cells bearing the 48 bp proto-telomere and the I-SceI expression cassette were treated with ahTET and aliquots were taken either prior to treatment (0 h) or after treatment (1 to 20 h). Genomic DNA was digested with ScaI and analyzed by Southern analysis using probes to *ura4*^+^ or *hph*^+^ (denoted by red bars above each locus). The I-SceI site is marked by a red triangle. The proto-telomere fragment is rapidly converted to the smaller *ura4*^+^ and *hph*^+^ fragments. The cleaved *ura4*^+^ and *hph*^+^ ScaI-I-SceI bands are indicated by partial ideograms of the original diagram of the proto-telomere. Molecular weight standards in kb are shown (M). The numbers in blue on top of the blot represent the hours after the induction. As these cells double every 4.5 hours, the 4 h time point is less than 1 population doubling. At the 20 h time point, the cells had doubled 3 times before growth stopped in stationary phase. As a control for loading, the blots were re-probed with a control *abo1*^+^ probe, as shown at the bottom. **(B)** Cells bearing the 0 bp proto-telomere cassette were treated and analyzed as in panel A. **(C)** Serial five-fold dilutions of cells bearing I-SceI sites at *lys1*^+^ and the 48 and 0 bp proto-telomere cassettes were spotted onto minimal media that lacks or has 9 µM ahTET. “*g*^+^” represents *gal1*^+^, while “*u*^+^” represents *ura4*^+^. **(D)** Quantitative analysis of survival of the strains in C after induction of I-SceI. Survival of both the 0 bp and 48 bp proto-telomere strains were significantly different than the strain bearing the I-SceI site at *lys1*^+^ (*p*<0.01, t-test). The 0 bp and 48 bp strains were not significantly different (*p*=0.09, t-test). Error bars show SEM from duplicate assays.

**Figure 4.**
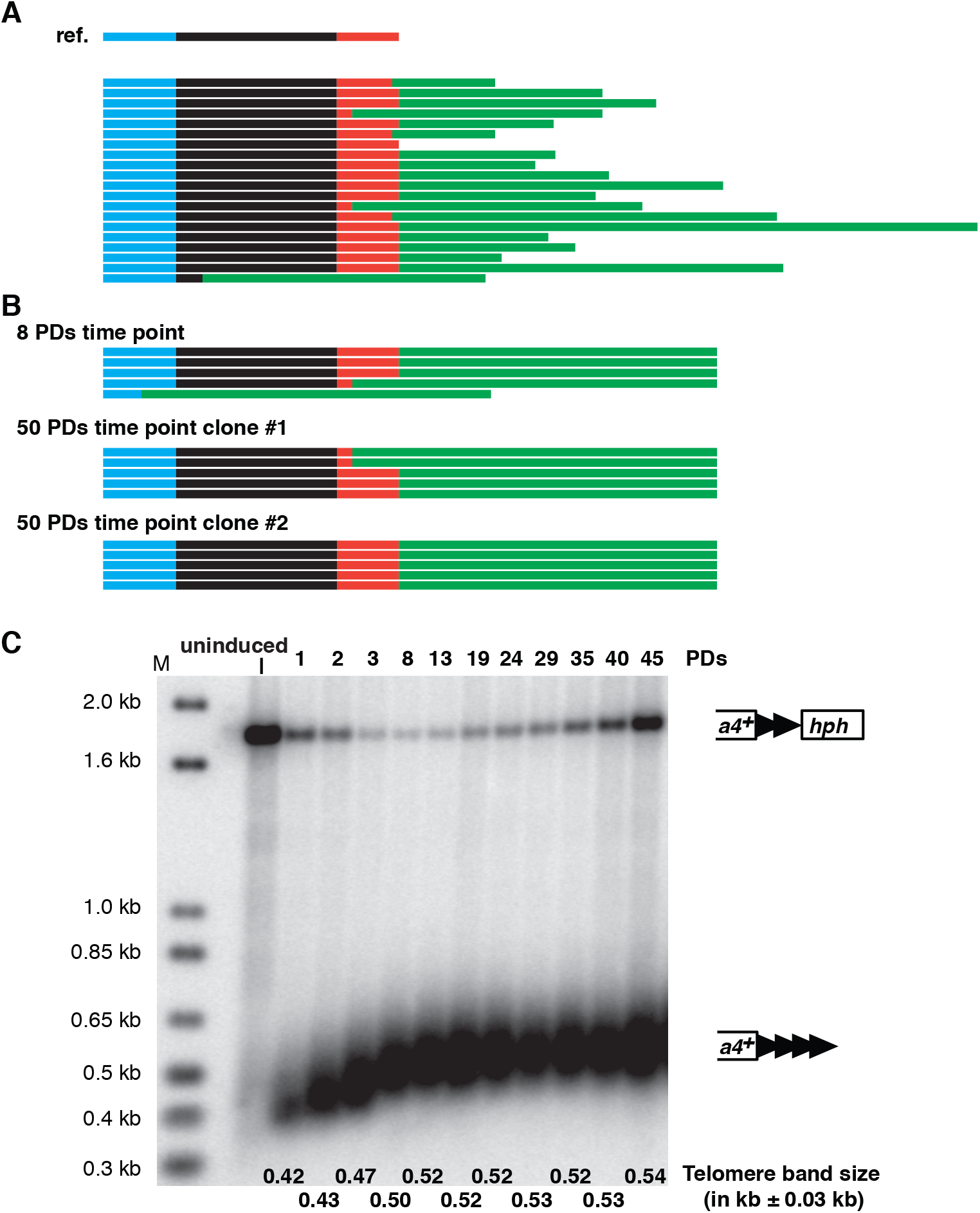
Analysis of the newly added telomere repeats at 48 bp proto-telomere. **(A) The sequence of newly added telomere repeats from cells at the 1 PD time point (4 h post-induction).** The top row (labeled “ref.”) shows the reference sequence of 48 bp proto-telomere in different colored bars. The bar in blue represents part of the 48 bp of telomere repeats. The black shows the polylinker sequence, and the red shows the I-SceI site including the overhang after I-SceI cleavage. The bars below are from 20 individual clones collected at the 1 PD time point. The green bar shows the newly added telomere repeats. Interestingly, all but one of telomere repeat addition was to the I-SceI site or polylinker sequences, similar to telomerase-mediated repeat addition in *S. cerevisiae* (54, 55, 84) and mammalian cells (38, 56). **(B) Telomere sequences cloned from the 8 PDs time point or different clones from the 50 PDs time point.** These fully elongated telomeres still retain the polylinker and I-SceI site in all but one case, indicating this conformation forms a stable telomere. Only the telomere repeat sequences closest to the addition site were shown. The detailed sequences in panel (A) and (B) are shown in Figure S1. **(C) Telomere repeat tracts are fully elongated by ~8 PDs after proto-telomere cleavage**. After induction of I-SceI, cells were grown for multiple PDs in liquid culture with 9 µM ahTET by serial dilution, and samples from different time points were processed for Southern blotting using *ura4*^+^ as probe as in Figure 3A. These data reveal that cells with an uncleaved proto-telomere had a growth advantage over cells with the new telomere, such that the cells with the uncleaved proto-telomere increased in proportion as cell grew. The uncleaved proto-telomeres most likely resulted from cassettes that were cut and healed by a DNA repair event that eliminated the I-SceI site.

**Figure 5.**
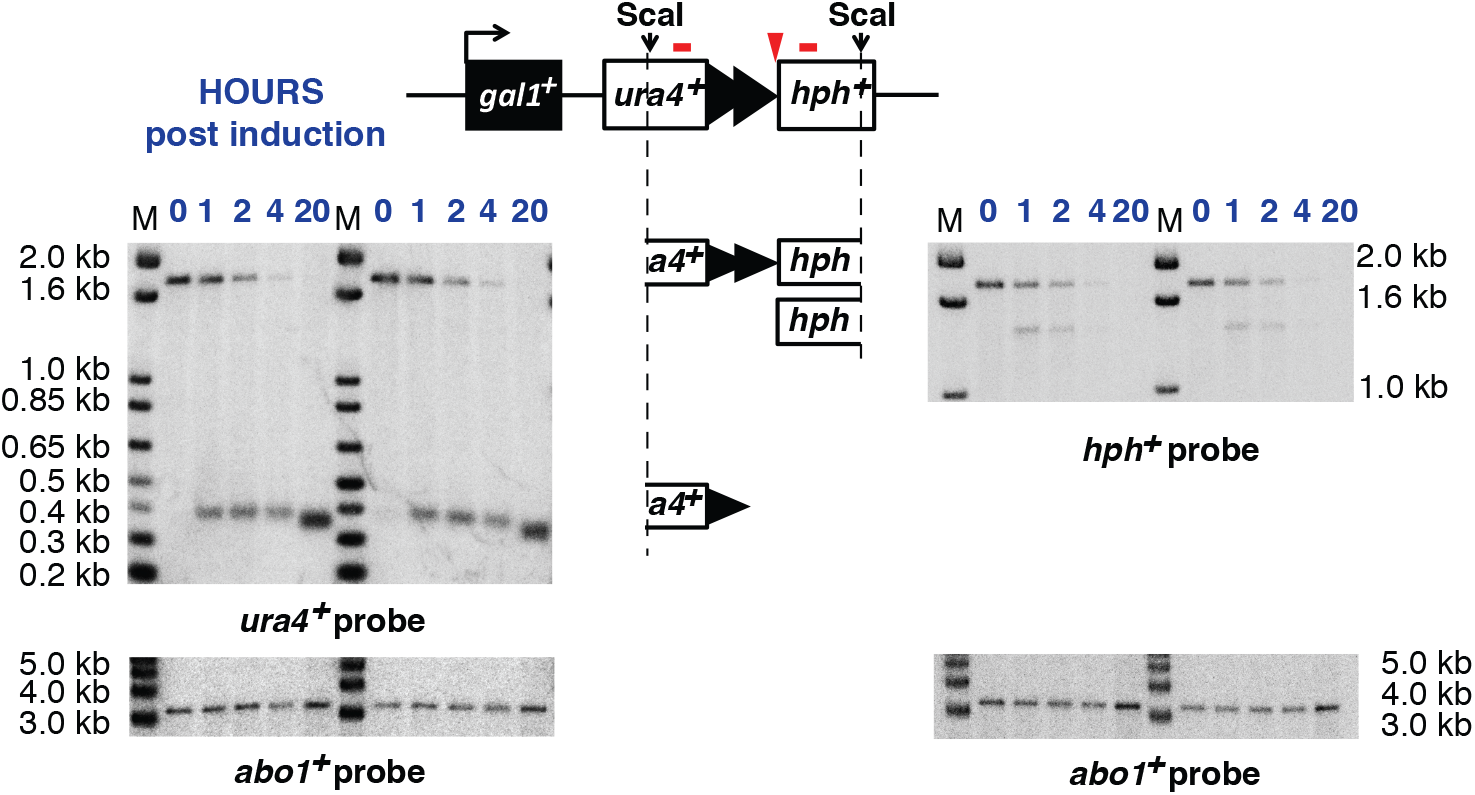
Telomerase null cells have no telomere elongation after induction of chromosome breakage. Two independent telomerase null (*ter1∆*) mutants were separately induced with ahTET and analyzed as in Figure 3A. The *ura4*^+^ and *hph*^+^ ScaI bands are indicated by partial ideograms of the uncut proto-telomere diagram.

The I-SceI-induced DSB at the 0 bp proto-telomere had a notably different fate from the 48 bp proto-telomere. The 0 bp proto-telomere strain displayed rapid cutting as demonstrated by the disappearance of the *ura4*^+^*-hph*^+^ fragment, and both I-SceI-generated terminal fragments were rapidly degraded (Figure 3B). Thus, I-SceI cutting at this locus was efficient and both sides of the DSB at the 0 bp proto-telomere were unstable.

### The genomic organization of *S. pombe* allows the efficient healing of subtelomeric DSBs

Double-strand breaks cause growth arrest in *S. pombe*, *S. cerevisiae*, human cells and other model systems while telomeres do not ((34, 50), reviewed in (39)). We therefore tested the effect of I-SceI cleavage at centromeric *lys1*^+^ and the subtelomeric 48 bp and 0 bp proto-telomeres. As expected, cleavage at *lys1*^+^ greatly impaired growth (Figure 3C and D). In contrast, cleavage at the 48 bp proto-telomere, which formed a telomere and lost subtelomeric repeated sequences, showed no detectable growth inhibition. Surprisingly, cells containing the 0 bp proto-telomere cassette also showed very little growth inhibition, with ~100% of the cells surviving (Figure 3C and D), even though the *ura4*^+^ fragment had been degraded in these cells (Figure 3B). The mechanism allowing this survival was unclear, because the DSB occurred in unique sequence, not in the sequences repeated in four telomeres.

To determine what process allowed the efficient growth of cells bearing the DSB formed at the 0 bp proto-telomere, we determined the chromosomal structure of three independent survivors. Phenotypic and genomic characterization revealed that the survivors had lost the *hph*^+^ gene and almost 19 kb of DNA internal to the I-SceI cleavage site. The degradation endpoint retained the *DUF999 protein family 7* gene (*DUF999-7*), a member of a gene family near the telomeres of chromosomes I and II, in which all genes are transcribed toward the centromere (Figure 6A). We hypothesized that nucleolytic degradation from the I-SceI site to the *DUF999 protein family 7* gene would allow recombination between gene family members to add a functional chromosomal end to IIR (Figure 6B), as recombination between repeats is known to be efficient enough to account for this high level of survival (58). To test this hypothesis, we determined the sequences adjacent to *DUF999-7* in the survivor strains and found sequences indicating recombination with *DUF999-8* on IIR or *DUF999-6* on IIL (Figure 6B and C). As the sequences from the *DUF999-8* and *DUF999-6* genes to their respective telomeres were almost identical (51, 52), the specific telomere captured by the DSB was not determined. Therefore, the *S. pombe* genome is structured to rapidly and efficiently heal DSBs near subtelomeres and maintain cell viability.

**Figure 6.**
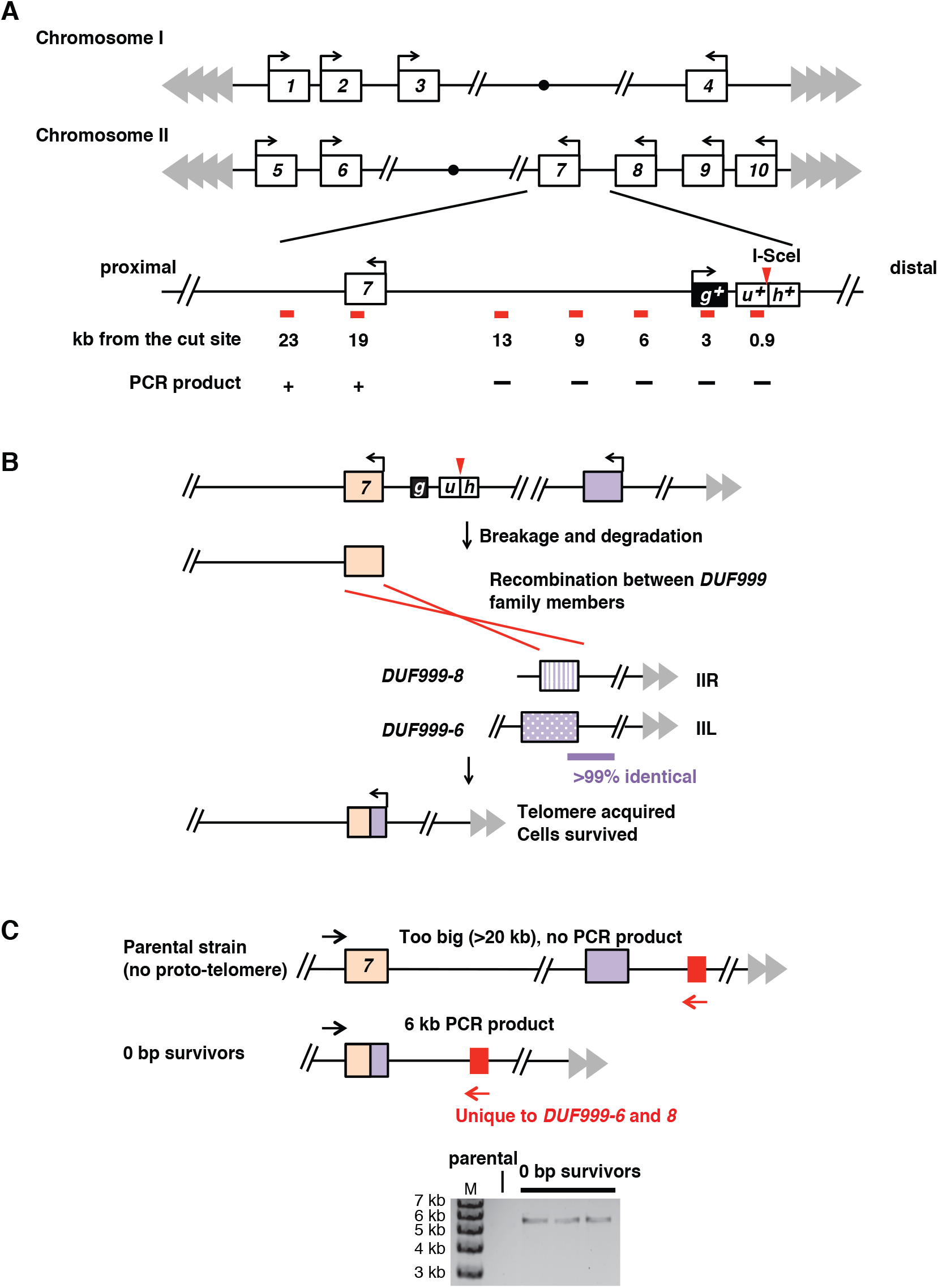
The genome organization of *S. pombe* allows efficient healing of the 0 bp proto-telomere. **(A) A map of the *DUF999* gene family on chromosomes I and II.** All 10 genes from this family are in the same orientation with transcription towards the centromere (the filled black circle). The chromosome III subtelomeres consist of scores of ribosomal RNA gene repeats (44), and are not shown. The region near *DUF999 protein 7* gene that is just internal to the 0 bp proto-telomere insertion site is expanded, showing the relative position of the proto-telomere and the distance of different primer pairs (shown as red bars) from the I-SceI site. A “+” indicates that a PCR product was obtained from each of the three surviving cleaved 0 bp proto-telomere strains tested and a “–” indicates that a product was not obtained. The *DUF999 protein 7* gene was the closest gene to the degradation endpoint. **(B) A hypothesis to explain how the *DUF999* gene family can provide a backup mechanism to rescue a DSB near the subtelomere.** After induction of a DSB at the 0 bp proto-telomere, DNA is degraded at both ends (Figure 3B). The generation of degraded DNA in the *DUF999 family protein 7* gene can produce a recombinogenic DSB that can undergo recombination with other *DUF999* genes (purple box) to acquire a new telomere. To test this hypothesis, we performed inverse circle PCR (see Materials and Methods) and determined the sequences that had been fused to the *DUF999 protein family 7* gene. We found a recombination donor that could be from *DUF999 protein family 3, 6* or *8*. **(C) PCR to confirm the recombination event.** *DUF999 family proteins 6* and *8* have a unique region (red box) that is absent from the *DUF999 family protein 3* gene region. PCR using a specific primer to this region (red arrow) and a unique primer at *DUF999 protein family* 7 (black arrow) revealed that the three strains that survived the induction of the DSB (the 0 bp survivors) were generated from the recombination between *DUF999 protein family 7* gene and *DUF999 protein family 6* or *8* genes. The sequence between *DUF999 protein family 6* and its telomere is nearly identical to the sequence between *DUF999 protein family 8* and its telomere, and thus the recombination event that rescued the DSB was not pursued further.

### Telomere formation initiates silencing of *ura4*^+^

To test silencing at a newly formed telomere, cells were assessed for expression of the *ura4*^+^ marker. Cells in which transcription of *ura4*^+^ is silenced, such as placing *ura4*^+^ near a newly formed telomere, are unable to grow on media lacking uracil (21, 23, 59). Cells were therefore induced for I-SceI expression overnight prior to plating on rich and selective media (Figure 7A). After induction, strains with the 48 bp proto-telomere grew poorly on medium containing hygromycin, indicating loss of the 3’ *hph*^+^ fragment (Figure 3A), or medium lacking uracil. However, the *ura4*^+^ gene was still present and could be amplified from these Ura^−^ Hyg^S^ colonies (Figure 7B). Amplification and sequencing of the PCR product containing the entire *ura4*^+^ gene revealed a wild type sequence. Therefore, establishing a new telomere silenced expression of the adjacent *ura4*^+^ gene.

**Figure 7.**
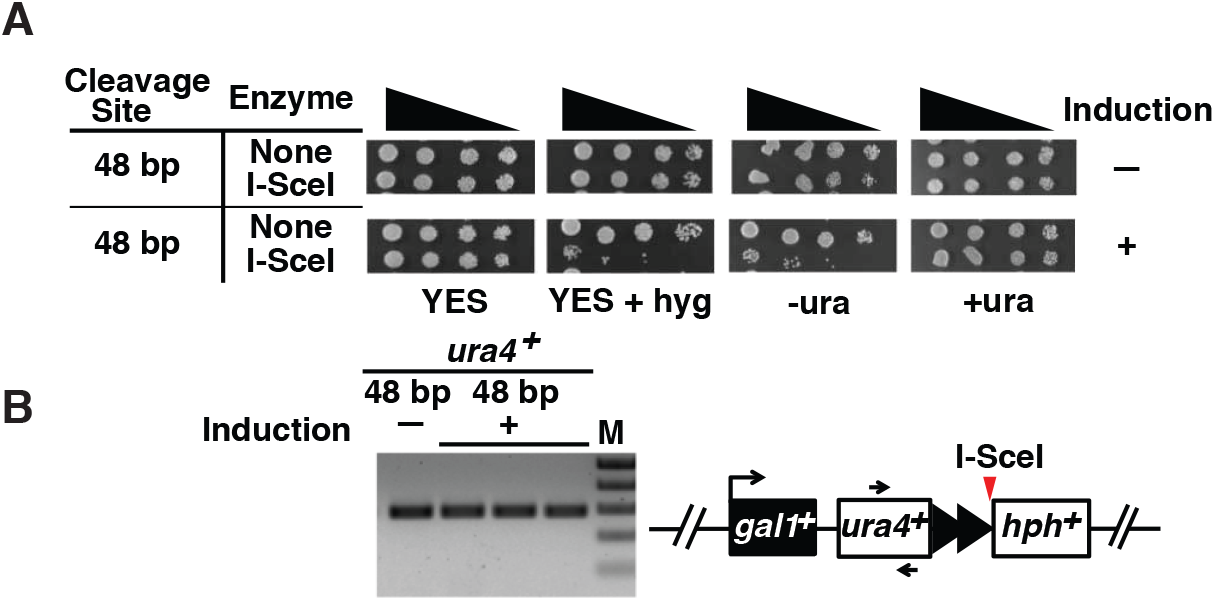
Telomeric *ura4*^+^ is silenced. **(A)** Growth of cells containing the 48 bp proto-telomere before and after 20 h of I-SceI induction was assessed by spotting five-fold serial dilutions of cells onto rich media plates (YES (71)) with and without hygromycin (hyg), or synthetic medium with (+ura) or without (-ura) uracil. **(B)** The presence of the *ura4*^+^ gene in untreated 48 bp proto-telomere cells prior to induction (–) or in three independent Ura^−^ Hyg^S^ colonies derived from ahTET treated cells (+) was tested by PCR. Primers are indicated by black arrows.

### The H3K9me2 heterochromatin mark spreads gradually after telomere formation, and is highly variable at full length telomeres

To determine if *ura4*^+^ silencing was due to heterochromatin formation, we tested whether levels of the heterochromatin specific histone modification, H3K9me2, were altered near the established telomere by ChIP-qPCR. H3K9me2 levels were determined using primers that amplify *ura4*^+^, *gal1*^+^ or chromosomal loci internal to the proto-telomere at varying distances up to 93 kb from the break (Figure 8A, red bars). We found that cells containing the uncut 48 bp proto-telomere had a localized peak of H3K9me2 near the insertion site, while the fully formed telomere showed a large increase of H3K9me2 spreading (Figure 8A). Spreading of the H3K9me2 mark was under nutritional control, as more spreading was observed in cells grown in rich medium than synthetic medium, even though telomere size was nearly identical under both conditions (Figure 8B). Therefore, similar to changes in *Drosophila* position effect variegation that respond to temperature (60) and the reversible silencing of *S. pombe* subtelomere-adjacent genes that are expressed in sporulation medium (25), heterochromatin domains in *S. pombe* also respond to environmental conditions. In cells with the 0 bp proto-telomere and no I-SceI gene, no such enrichment of H3K9me2 was found (Figure 8C). The localized H3K9me2 peak in the uninduced cells with the 48 bp proto-telomere did not spread into the distal end (Figure 8D).

**Figure 8.**
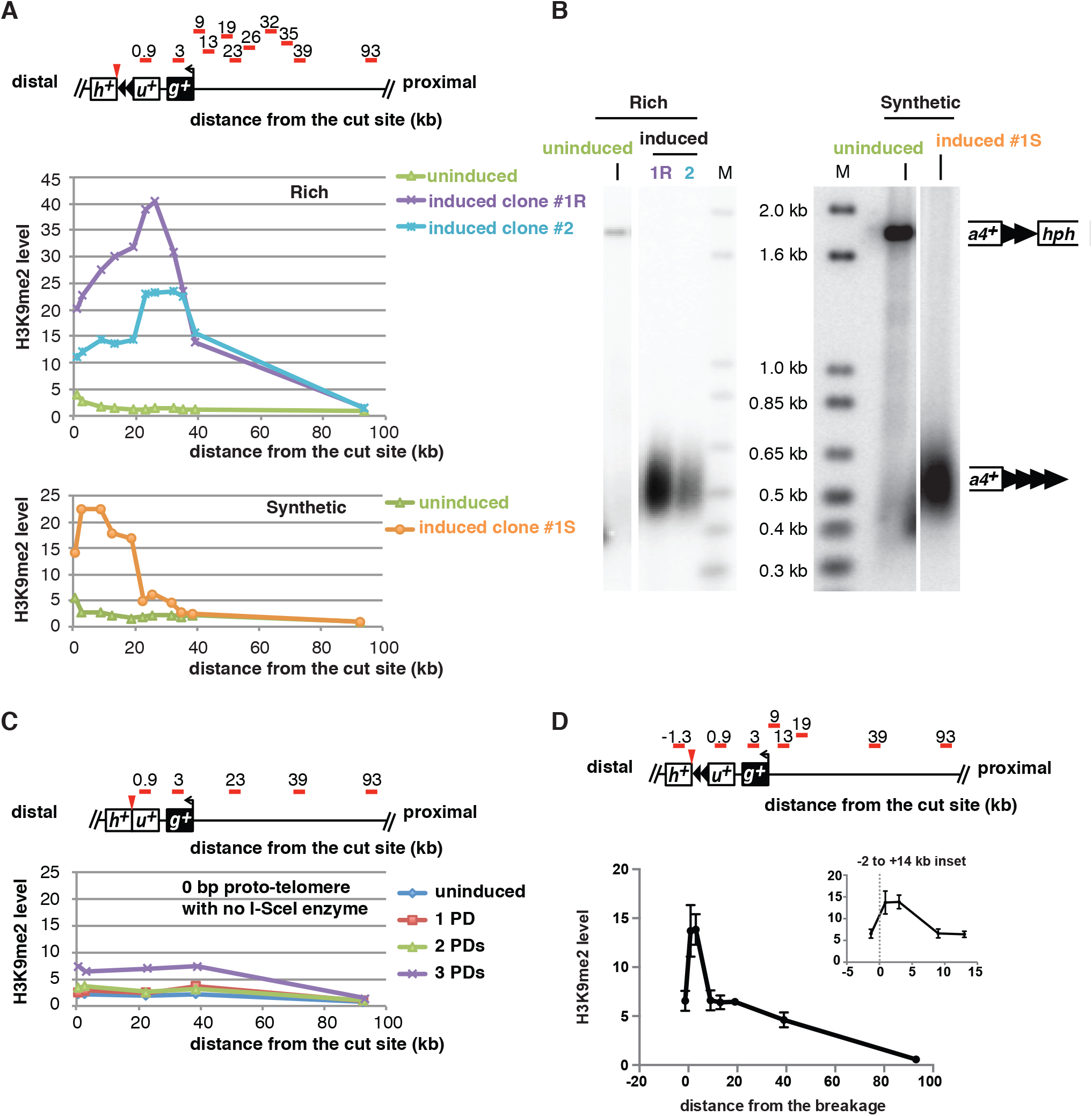
Different telomeric H3K9me2 domains form in cells with telomeres of similar size. **(A) The level of histone H3K9me2 enrichment at each locus (primers locations are shown as red bars) is shown.** The purple and cyan lines (top) show two individual clones of 48 bp proto-telomere grown in rich medium (YES + 3% glucose) for 50 PDs after induction, which both peak at the 26 kb locus. In contrast, cells grown in synthetic medium (EMMG with uracil + 2% glucose)(bottom) have the highest H3K9me2 level at a locus much closer to the newly formed telomere. The green line (both top and bottom) represents the uninduced 48 bp proto-telomere, which shows a small increase next to the telomere repeat tracts in the proto-telomere. **(B) The established telomeres have similar lengths in cells from rich (left) or synthetic media (right).** The lanes with DNA from cells in panel A: uninduced cells are labeled in green while those with induced cell DNAs are in purple or cyan (left, in rich media) or orange (right, in synthetic media). Southern analysis used the *ura4*^+^ probe, as in Figure 3A. Molecular weight standards are labeled with “M”. **(C) 0 bp control cells do not have increased H3K9me2 level after induction.** The levels of histone H3K9me2 enrichment at each locus are shown. Distances are relative to the I-SceI cut site. Red bars indicate the PCR probes. A schematic of the 48 bp proto-telomere shows the location of the 0.9 and 3 kb probes in *ura4*^+^ and *gal1*^+^, respectively. No major enrichment was seen in cells with 0 bp proto-telomere and no I-SceI gene. **(D) The H3K9me2 peak is localized on telomeric repeats in uninduced cells.** The levels of histone H3K9me2 enrichment at each locus are shown. Distances are relative to the I-SceI cut site. Red bars indicate the PCR probes. The leftmost -1.3 kb probe recognizes the *hph*^+^ coding sequence and is 1.3 kb from the I-SceI site. The average and range of two independent tests are shown. The inset shows the H3K9me2 level from -2 to +14 kb and the I-SceI site at 0 kb is marked by a dashed grey line.

To understand the relationship between the formation of a functional telomere and the establishment of the telomeric heterochromatin domain, we performed a kinetic analysis of H3K9me2 levels while the new telomere was forming. Upon induction of telomere formation, heterochromatin spreading was monitored in cells grown continuously for 8 PDs. To examine cells at longer time points in the absence of cells that have healed the I-SceI cut to retain the *hph*^+^ fragment and subtelomere (Figure 4C), cells from PD 2 were used to isolate single, Hyg^S^ colonies that were subsequently cultured and analyzed (Figure 9A).

**Figure 9.**
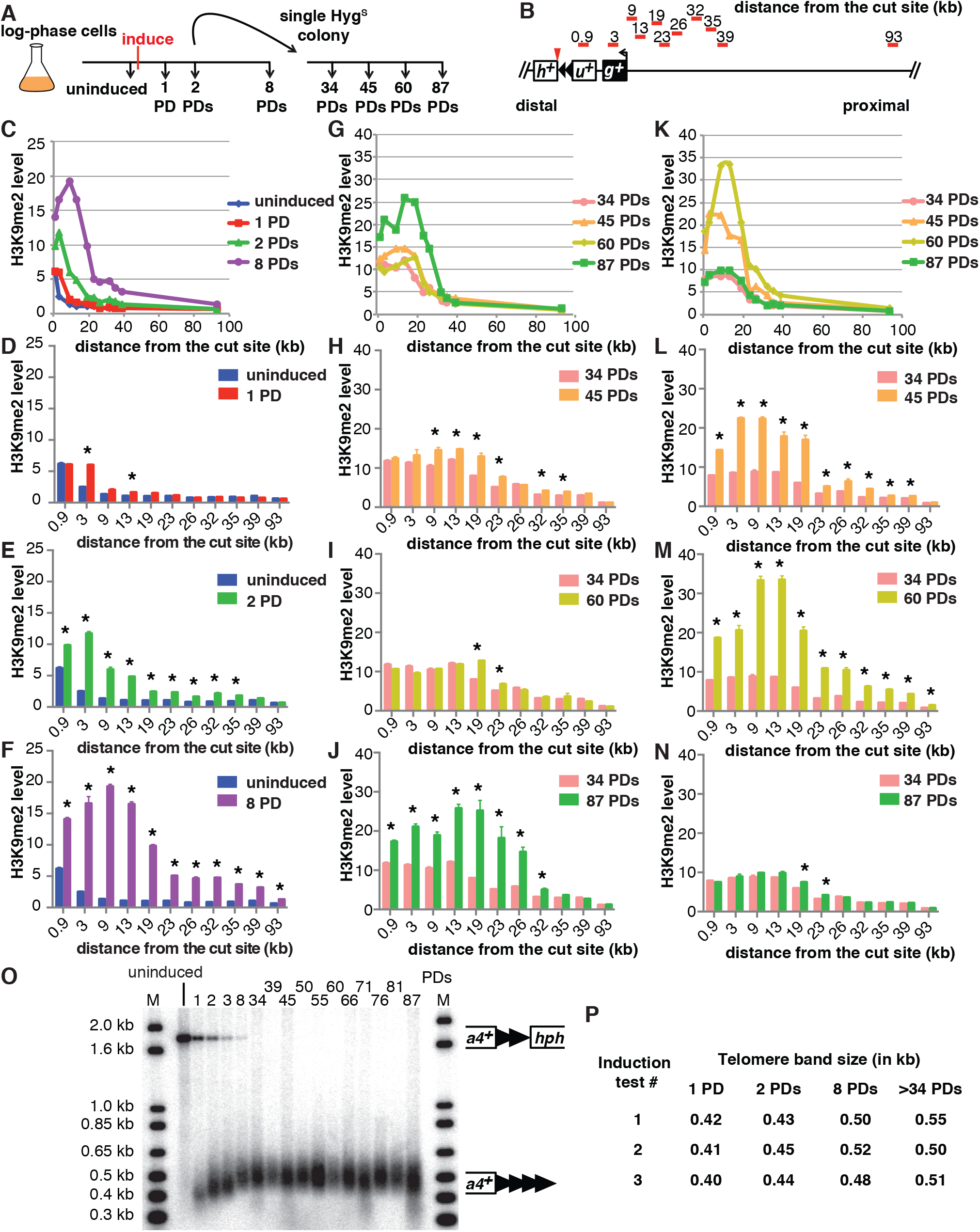
The new H3K9me2 domain forms gradually as the new telomere reaches equilibrium. **(A)** Telomere formation is induced and samples were taken at different time points for analysis of telomere length and H3K9me2 levels. **(B)** The primer sets used to monitor the levels of histone H3K9me2 enrichment at several loci are shown as red bars. Distances are relative to the site of I-SceI cut site, represented as a red triangle. The 0.9 and 3 kb probes are in *ura4*^+^ (*u*^+^) and *gal1*^+^ (*g*^+^), respectively. “*h*^+^” is *hph*^+^. **(C)** Kinetic analysis of the levels of histone H3K9me2 enrichment at several loci over 8 PDs, normalized to total histone H3 levels at each locus (Materials and Methods), are shown. **(D-F)** H3K9me2 levels at each locus and time point compared to the levels in uninduced cells bearing the 48 bp proto-telomere. Statistically significant differences for each locus compared to the uninduced level (*p*<0.05, t-test) are marked by an asterisk. Error bars show the SEM of triplicate assays. **(G)** Kinetic analysis of heterochromatin spreading of a single Hyg^S^ colony (panel A), shown as in C. **(H-J)** H3K9me2 levels at each locus and time point compared to the 34 PDs culture for the Hyg^S^ colony shown in G. Statistically significant differences (*p*<0.05, t-test) are indicated by an asterisk. **(K)** Kinetic analysis of a second independent Hyg^S^ colony, as in C and G. PD 0-8 of this experiment are shown in Figure S2A. **(L-N)** Analysis of individual time points shown in K as in H-J. **(O)** Telomere size was measured at different time points after induction by Southern analysis using a *ura4*^+^ probe as in Figure 3A. Molecular weight standards are labeled with “M”. **(P) Telomere elongation is nearly identical in independent telomere formation experiments.** The modal terminal restriction fragment (TRF) sizes (the *ura4*^+^-telomere repeats band) of the newly formed telomere at early and late population doublings (PDs) after induction with ahTET, were determined as the most intensely hybridizing part of the band on Southern blots. Band sizes on these blots vary by approximately ± 0.03 kb. Induction #1 sizes are from the formation experiment shown in panel K-N & Figure S2A. Induction #2 sizes are from the formation experiment in panel O. Induction #3 sizes are from the formation experiment in panel C-J.

From 0 to 8 PDs, the H3K9me2 level gradually increased and peaked at 9 kb from the cut site (Figure 9C-F and Figure S2A-C) and telomeres were elongated gradually (Figure 3A and 9O). At the 1 PD time point, cells had short functional telomeres as the *ura4*^+^ telomeric fragment was stable and slightly elongated (Figure 3A and 9O), but H3K9me2 level barely increased (Figure 9C and D). At 2 and 8 PDs, the size of heterochromatin slowly increased toward the centromere (Figure 9C, E and F) and telomere length reached its equilibrium state at 8 PDs (Figure 9O).

Surprisingly, from PDs 34 to 87, independently formed telomeres from four similar induction assays showed differences in the amount of heterochromatin at different times in the presence of constant telomere length. These experiments showed spreading of the level of H3K9me2 to a domain of similar size (Figure 9G and K), and H3K9me2 levels were very similar at the most internal loci at all time points. However, one line (Figure 9G-J) showed a peak of heterochromatin at 19 kb from the new telomere that was nearly constant from 34 to 60 PDs, followed by a significant increase by 87 PDs. A second line showed an increase at 60 PDs that was maintained at 87 PDs (Figure S2D). In contrast, the remaining two lines (Figure 9K-N and S2E) showed a similar internal peak that increased from 34 to 60 PDs, followed by a significant decrease by 87 PDs. Southern analysis revealed that telomere lengths in different formation experiments were indistinguishable at all of these time points (Figure 9O and P). Therefore, spreading of the telomeric H3K9me2 mark was dynamic, even though the telomere maintained a constant repeat tract size during this time.

## Discussion

We have constructed the first inducible *S. pombe* telomere formation system, and used it to show that telomeric regions have unexpected properties in healing DSBs and in the kinetics of heterochromatin domain formation and spreading. While inducing a DSB near the middle of the chromosome arm caused a significant growth inhibition, DSBs in the subtelomeric region at the 0 bp or 48 bp proto-telomere did not (Figure 3C and D). The 0 bp proto-telomere lacking any telomere repeats showed DNA degradation on both sides of the DSB (Figure 3B), and revealed a backup mechanism to restore telomere function by recombination between a family of subtelomeric repetitive elements (Figure 6). In contrast, the 48 bp proto-telomere with telomere repeats on the centromeric-side of the DSB was stable and a substrate for telomere repeat addition, behavior identical to a functional short telomere (33, 52, 61)(Figure 3A, 4 and 9O). Even though formation of a functional telomere was rapid, establishing the telomere-dependent heterochromatin domain was much slower. The H3K9me2 domain spreads gradually from 0 to 8 PDs, when telomere length reaches equilibrium state. The slow spreading of heterochromatin is consistent with the facts that the essential telomere functions in chromosome stability are independent of heterochromatin (22, 57), and that this heterochromatin domain is a secondary consequence of telomere formation. The slow spread of the heterochromatin domain raises the possibility that chromatin domain formation in other biological contexts (e.g. metazoan development, tumorigenesis, senescence) also requires several cell divisions. After telomere repeat tracts reach their final length, the size of the H3K9me2 domain remains dynamic, indicating that this extended spreading is independent of telomere length.

The backup mechanism to rescue telomere function in response to a subtelomeric DSB most likely reflects the similarities between the genome structure of *S. pombe* and metazoans. The three nuclear chromosomes of *S. pombe* have complex subtelomeric regions (51), with repeats oriented in a way that allowed recombination to attach or copy a functional telomere to the broken chromosome end (Figure 6). Mammalian genomes also contain a large number of repeats, and small deletions at the border between telomeric euchromatin and heterochromatin are unlikely to cause a phenotype in diploid cells, in contrast to large telomeric deletions that have developmental consequences (61–63). The *S. pombe* results therefore suggest that a direct examination of induced DSBs near the border of the mammalian heterochromatic subtelomere may reveal a similar mechanism for rescuing telomere function.

The newly formed *S. pombe* telomere revealed an unusual heterochromatin domain compared to the uncleaved 48 bp proto-telomeres. The uncleaved 48 bp proto-telomere showed only a peak of H3K9me2 levels that were centered at the 48 bp telomere repeats (uninduced in Figure 8 and 9C), consistent with the internal telomeric repeats initiating low levels of silencing (22, 23). In contrast, the established telomere with a functional chromosome end formed an internal heterochromatin domain that peaked from 9 to 26 kb from the telomere (Figure 9C, G, K). The reason for this internal peak, as opposed to a peak immediately adjacent to the telomere repeats, is unknown. The location of this internal peak was different in cells grown in rich media versus synthetic media (Figure 8A), suggesting that the peak location is not completely sequence dependent. This *S. pombe* telomere formation system will thus provide a useful tool for future studies to examine the *cis-* and *trans-*acting factors that regulate the positioning of this heterochromatin peak.

The telomere formation system revealed a slow and dynamic spreading of the telomeric heterochromatin domain that was not predicted by previous studies. In recent work where a synthetic *S. pombe* heterochromatin domain was established by conditionally tethering the H3K9 methyltransferase to an expressed gene, release of the tethered methyltransferase caused the H3K9me2 mark to be lost a few hours later (~1-2 PDs)(12, 13), much faster than the 8 PDs (48 h) required to form the internal heterochromatin peak (Figure 9C). Assembly of a transcriptionally silenced chromatin in *S. cerevisiae*, which does not involve H3K9me2, has been monitored and also forms at a much faster rate. Overexpression of a silencing protein unique to the budding yeast, Sir3, in wild type cells extended existing Sir3-containing chromatin domains (30–32). An independent approach that used chemical inhibition of silencing and followed its establishment after the inhibitor was withdrawn (64) showed that Sir3 silent chromatin was significantly extended or re-established in these studies within ~4 hours (~1-2 PDs). Complete modification of the histones as a consequence of Sir3 spreading, however, did require additional population doublings (30). In contrast to these events in yeasts, formation of a synthetic heterochromatin domain in murine cells from a tethered silencing factor took much longer (~5 days) to form the steady-state 10 kb heterochromatin domain (65). While the slower kinetics could be a consequence of the murine tethering system, the *S. pombe* telomere formation results suggest that the assembly of H3K9me2-dependent heterochromatin domains is an intrinsically slower process compared to its disassembly.

The telomeric H3K9me2 chromatin domain formed in two distinct phases following telomere formation in a wild type strain background. The first was the spreading over 8 PDs to form the domain where H3K9me2 peaks near 9 kb from the telomere (Figure 10), even though a substantial fraction of the telomeres were already normal length by 2 PDs (Figure 9O). This mechanism is consistent with current models of spreading (24, 40), in which the extension of the H3K9me2 modification from its nucleation site (e.g. the newly formed telomere) can only occur in S-phase after DNA replication when new chromatin is formed, then S-phase exit may limit the extent of heterochromatin extension for that cycle. The second phase occurs during continued propagation of cells with fully elongated telomeres, where the internal peak of H3K9me2 chromatin from 9 to 26 kb can significantly increase or decrease. How these changes occur is unknown, and could reflect methylation of the Lys9 residues on both H3 amino termini in the histone octamer or the presence of a subpopulation of cells in the culture that had not formed this heterochromatin domain. The internal peak of H3K9me2 chromatin may be indicative of a cryptic enhancer of H3K9me2 modification that is activated by this modification spreading from the telomere. In these hypotheses, the changes in H3K9me2 levels occur slowly in continuously growing cultures, consistent with the assembly of new chromatin every new S-phase (Figure 10), although the active replacement of nucleosomes by chromatin remodeling factors outside of S-phase cannot be ruled out.

**Figure 10.**
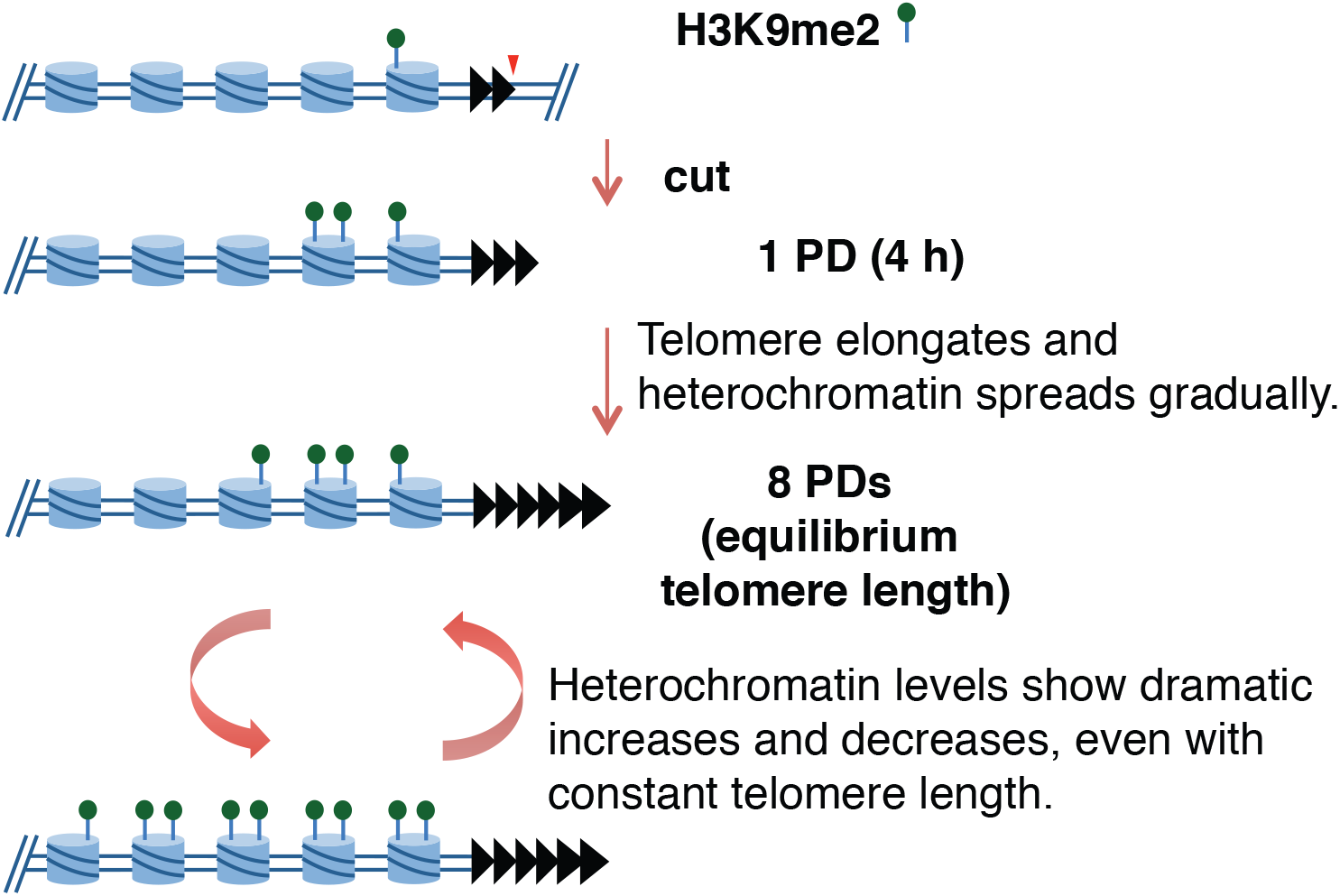
Hypothesis for heterochromatin formation over multiple cell divisions. The 48 bp proto-telomere (black triangles) has a low level of H3K9me2 in nucleosomes (blue cylinders) of the adjacent loci. While telomeres are immediately functional for end protection upon I-SceI breakage, only a small amount of H3K9me2 mark has been established by 1 PD. Spreading gradually increases over 8 PDs. However, the amount of spreading still varies as cells continue to grow, even though telomere repeat tract length is constant. These increases may reflect more cells where both N-termini of histone H3 are modified by lysine 9 dimethylation, a larger fraction of cells in the culture that have this modification at these loci or both.

Recently, Obersriebnig *et al.* examined the spreading of gene silencing and the H3K9me2 modification in the *S. pombe* silent mating type region after reintroduction of the *clr4*^*+*^ gene into a *clr4Δ* cell by mating. Silencing at different distances from the initiation site was monitored by following the loss of expression of fluorescent protein genes for the first several generations, and H3K9me2 spreading was followed ~30 divisions later by ChIP (14). Numerical modeling of the rates of spreading, assuming that fluorescent protein gene silencing was due to H3K9me2 spreading, could be divided into global and local effects that produced outcomes similar to their experimental observations. Spreading in the first few cell divisions was consistent with a linear spreading from the initiator in the silent mating type region (the *cenH* repeat), similar to spreading of the H3K9me2 mark from our newly formed telomere. Despite the similarities of the two systems in these initial stages of spreading, it is important to note that the silent mating type region is a highly specialized and well-studied structure containing initiators, enhancers and boundary elements that both promote and confine heterochromatin to a defined region to permanently extinguish gene expression as an essential part of the fission yeast life cycle (66–68). The newly formed telomere does not contain any known elements of this type, and may be more similar to the heterochromatin domain formation that occurs during development or senescence that encompasses genes that are expressed in difference cell types (5–7, 25). Additional work on the newly formed telomeric H3K9me2 domain will therefore be required before a detailed comparison with the silent mating type region can be made.

The slow extension of the H3K9me2 modification from its nucleation site (e.g. the newly formed telomere) has functional consequences for the formation of larger chromatin domains, which may require many cell cycles. The rate of transition from one cellular state to another during development or aging would be slowed by formation of a chromatin domain. This rate may well be increased early in development as oocytes have large amounts of maternally deposited histones and histone modifying enzymes (69, 70), and the increased levels of chromatin components and modifying enzymes could increase the kinetics of chromatin domain formation. In somatic cells where the modifiers may be at lower levels, the kinetics of domain formation would be slower and may impede aging and tumorigenesis. The *S. pombe* telomere formation system will provide an ideal model for testing these ideas and identifying the rate-limiting components in chromatin domain formation with broader implications for metazoans.

## Materials and Methods

### Strains and Media

All *S. pombe* strains used in this study are shown in Supplementary file 3. Selection for strains containing telomere cassettes was performed in Edinburgh Minimal Media with sodium glutamate (EMMG) substituted for ammonium chloride without uracil and with appropriate amino acid supplements and 100 µg/ml Hygromycin B Gold (InvivoGen)(71). Non-selective growth of strains bearing the telomere cassettes was done in EMMG with uracil and other appropriate amino acid supplements and without hygromycin. Preparation of 10 mM anhydrotetracycline stock and plates was performed as in (34). 5-FOA plates are Yeast Nitrogen Base plates with 1 mg/ml 5-FOA (Toronto Research Chemicals, Inc.)(72) and with the appropriate supplements. All recombinant DNA procedures were carried out in NEB 5-alpha (New England Biolabs) and TOP10 (Life Technologies) competent cells.

### I-SceI Expression Vector

I-SceI is produced from a synthetic gene with optimized *S. pombe* codons (47) and expressed as a protein with two N-terminal SV40 nuclear localization signals (NLS) fused to I-SceI. I-SceI expression is under the Cauliflower Mosaic Virus 35S promoter (CaMV35Sp), which is regulated by the tetracycline repressor (TetR). The TetR protein is produced from the *adh1*^*+*^ promoter in the same cassette as I-SceI (73). pFA-LEU2-I-SceI was produced by a 5-part recombination cloning in *S. cerevisiae*, rescued to bacteria, and verified by DNA sequencing (74). An I-SceI site on the vector backbone was removed by site-directed mutagenesis. Additional cloning details are available upon request. The vector and its sequence have been deposited with Addgene.

### Telomere Cassette

The most terminal unique region of *S. pombe* Chr IIR was found to be the 2 kb region 3’ of the *gal1*^+^ gene 3’-UTR (44). The proto-telomere cassette containing *ura4*^+^, 0 or 48 bp of telomere seeding sequence, and the *hph*^+^ gene encoding hygromycin resistance was constructed in the vector pRS315 by a 5-part recombination cloning in *S. cerevisiae*. The junctions between DNA fragments were verified by colony PCR and the plasmids were rescued to bacteria and sequenced. Additional cloning details are available upon request. The vector and its sequence have been deposited with Addgene.

### Construction of the *I-SceI*-*lys1*^+^ Allele

The I-PpoI site in the plasmid pSS23 (34) was replaced by the I-SceI site by standard cloning. Transformation, selection for the hygromycin resistance gene *hph*^+^, and confirmation of integration of I-SceI at *lys1*^+^ in *S. pombe* was done as before (34).

### Induction of I-SceI

Cells containing the telomere cassettes were grown under selection overnight and diluted to a final volume of 230 ml at 5.5 x 10^6^ cells/ml in non-selective media and grown for 3.75 hours. Untreated cells (3-5 x 10^8^) were removed and pelleted, washed once with sterile water and frozen at -80°C. Anhydrotetracycline (ahTET) was then added to a final concentration of 9 µM. Cells then were collected at various time points and pelleted, washed, and frozen as above. Genomic DNA was extracted (75) from the frozen pellets for Southern analysis as below. I-SceI cleavage at *lys1*^+^ was performed and analyzed as in (34), except ahTET was added at a final concentration of 9 µM.

### Southern Blot Analysis

Cells (3-5 x 10^8^) were collected at each time point and used to prepare genomic DNA (75). Genomic DNA (5 µg) was digested with 20 units of ScaI and analyzed via Southern blot with ^32^P-labeled probes produced by PCR with u4ScaProbe_S + u4ScaProbe_AS or SV40ScaProbe_S + SV40ScaProbe_AS (Supplementary file 4). Purified PCR product (50 ng) was denatured and treated with 10 units of Klenow (New England Biolabs) in the presence of primers (final concentration 0.25 µM) and 100 µCi of alpha-^32^P-dATP (3000-6000 Ci/mmol, PerkinElmer) in a 40 µl reaction at 37°C for 30 min. The probe was purified in a G-25 spin column and 2-10 x 10^6^ counts per minute (cpm) was used in Southern blot hybridization. Pre-hybridization and hybridization performed with PerfectHyb (Sigma) as described (76). Stripping of membrane performed in buffer containing 0.5% SDS and 0.1x SSC and heated to 100°C for 2 × 15 minutes.

### Telomere PCR and Sequencing

Telomere PCR was performed as previously described (76, 77) with primers u4-teloPCR-1S and BamHI-G_18_ (Supplementary file 4) using genomic DNA from 1, 8 and 50 PD(s). Cells from 8 PDs (48 h) were struck for single colony and tested for hygromycin sensitivity. Two hygromycin-sensitive colonies were used for this analysis as 50 PDs clone #1 and #2. The purified PCR products were cloned into TOPO vector (Life Technologies) and sequenced using M13F or M13R primers (Supplementary file 4) at the Lerner Research Institute Genomics Core.

### ahTET Plating Assay

The spot test assay was performed by spotting 5-fold serial dilutions onto the indicated plates as in Sunder *et al*. (34). Strains containing proto-telomere constructs were grown without ahTET and under selection for the telomere cassette, and then plated on non-selective EMMG with and without ahTET. For the quantitative plating assay, cells were plated onto non-selective EMMG with or without ahTET at 300 cells per plate and grown for 7 days. The average number of colonies from three individual plates with ahTET was normalized to that from plates without ahTET for strains containing the I-SceI gene. This was then normalized to the same ratio of control cells without the I-SceI gene. Statistical comparisons were performed using GraphPad Prism version 6.0 (GraphPad Software).

### Selection of 0 bp Survivors

*S. pombe* cells containing the 0 bp proto-telomere were induced with ahTET (9 µM final concentration) and grown overnight in liquid EMMG. Cells were struck for single colonies on rich media and grown for 3 days. The resulting colonies were tested for sensitivity to hygromycin (100 µg/ml). DNA was extracted from 3 separate isolates that were sensitive to hygromycin and analyzed via PCR using primers listed in Supplementary file 4 to determine which sequences had been deleted after initiating the DSB at the 0 bp proto-telomere.

### Mapping of 0 bp Survivors

The recombination site was determined using inverse-circle PCR. Briefly, genomic DNA from 3 separate isolates (5 µg) was digested with 20 units of *EcoRI* for 16 h at 37°C followed by inactivation at 65°C for 20 min. A portion of the digestion (1 µg) was diluted in a ligation reaction to a total volume of 200 µl using 40 units of T4 DNA Ligase (New England Biolabs) for 16 h at 18°C. The ligation was ethanol precipitated and resuspended in 10 µl of 10 mM Tris-1 mM EDTA, pH 8.0. Half of the product was amplified with primers 07c-2-AS-rv&compl + BsrDI-map-AS and the product was sequenced with the same primers (Supplementary file 4). The resulting sequence was subjected to BLAST analysis and aligned to the *S. pombe* genome (44).

### Silencing Assay

Cells containing the telomere cassettes were grown under selection overnight. Cells were then transferred to 5 ml of non-selective media at a concentration of 5.5 x 10^5^ cells/ml and allowed to recover for 3.75 h before addition of ahTET (9 µM final concentration). Cells (1 x 10^6^) were collected before and after overnight induction with ahTET and 5 x 10^5^ cells were plated in five-fold serial dilutions on plates with the media indicated and grown for 3 or 4 days at 30°C.

### Analysis of *ura4*^+^ in the Ura^−^, 5-FOA Resistant Colonies

Single colonies resistant to 5-FOA (5-fluoro-orotic acid, which Ura4 converts to a poison) after induction of I-SceI were tested for hygromycin sensitivity on rich media. DNA was extracted and analyzed for the presence of *ura4*^+^ by PCR using 5 PRIME HotMaster *Taq* DNA Polymerase according to manufacturer's instructions and primers ura4ChIP_F + ura4ChIP_R (Supplementary file 4) and an extension time of 1.0 min for 25 cycles (MJ Research PTC 200 Thermal Cycler). A positive control for all PCRs was performed in parallel using primers SPBPB2B2.07c-ChIP-S + SPBPB2B2.07c-ChIP-AS (Supplementary file 4) to amplify the *DUF999* protein family 7 gene and produced a product in all reactions.

### ChIP Assay

Cells in 300 ml at 0.8-1.2 OD_600_ were cross-linked with 1% formaldehyde, and washed twice with cold HBS buffer (50 mM HEPES-NaOH pH 7.5, 140 mM NaCl). Cell pellets were stored at -80°C. For a saturated culture, cells were diluted to the above OD_600_ for cross-linking. At 2 PDs, cells were struck for single colony on rich media for 3-4 days. The resulting colonies were tested for sensitivity to hygromycin (100 µg/ml). A hygromycin-sensitive colony was inoculated in non-selective EMMG with ahTET and cells from serial dilutions were collected for analysis. All subsequence steps were performed at 4°C. Cell pellets were resuspended in ChIP-lysis buffer (78) and lysed using mechanical disruption by beads-beater (Bio Spec Mini-Beadbeater-16) with 0.5 mm glass beads (Biospec 11079105) using 4 cycles of 45 sec followed by 60 sec on ice. The lysate was sonicated for 10 cycles on maximum power (30 sec ON and 59 sec OFF) in a Diagenode Bioruptor XL with sample tubes soaked in an ice water bath. Solubilized chromatin protein (2-4 mg) was used for each ChIP while 5 µl was saved as Input. Antibodies (2 µg) against H3K9me2 (Abcam ab1220) or total H3 (Abcam ab1791) were added to lysate and incubated while rocking for 4 h at 4°C. Dynabeads Protein G (50 µl, Life Technologies) was then added to lysate for rocking overnight at 4°C. Beads were washed with ChIP lysis buffer, ChIP lysis buffer with 500 mM NaCl, Wash buffer and TE buffer (10 mM Tris, 1 mM EDTA pH 7.5) successively (78). Beads were then resuspended in 145 µl of TES (1 × TE with 1% SDS). Supernatant (120 µl) was recovered and incubated in a Thermomixer at 65°C, 1000 rpm overnight to reverse cross-linking. For Input samples, TES buffer (115 µl) was added and incubated in the Thermomixer with the ChIP samples. Samples were treated with RNaseA and ProteinaseK, and purified by QIAgen PCR purification column (79). All time points from the same induction assay were processed for ChIP assay at the same time.

### qPCR Analysis for ChIP

Input samples were diluted to 1/100 with ddH_2_O while beads-only-ChIP, H3-ChIP and H3K9me2-ChIP samples were diluted to 1/10. Template DNA (4 µl) were added to 5 µl of Roche LightCycler 480 SYBR Green I Master (2X) and primers were added to a final concentration of 0.6 µM for a 10 µl total reaction volume. Each sample was run in triplicate on the same 384-well PCR plate (Roche LightCycler 480 Multiwell Plate 384, clear) in a Roche LightCycler 480. Each ChIP assay was performed at least three times independently. H3K9me2 levels were normalized to the total H3 levels at each locus (80– 82), and each ratio was normalized to *act1*^+^ control locus in the same ChIP (83). Fold enrichments (*FE*) were calculated using the delta-delta-Cq method for each locus at each time point, as followed for a locus of interest (*loi*),

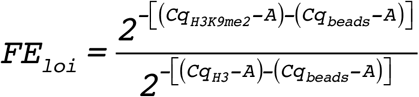

where

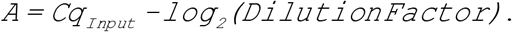

Then *FE* of the *loi* was normalized to *FE* of *act1*^+^ to generate the final Fold Enrichment of H3K9me2 at each locus.

## Acknowledgements

We would like to thank Dr. Steven Sanders for use of strains and materials, all the members of the Runge lab, and Drs. Bibo Li, Derek Taylor, Peter Harte and Hung-Ying Kao for discussion and critical comments on the manuscript. This work was supported by National Institutes of Health Grants R01GM050752 and R01AG019966 and National Science Foundation Grant 1516220 to KWR. KLB was supported by National Institutes of Health Grants R01HL055666 and R01HL081093.

## References

1. Chaligné R, Heard E. 2014. X-chromosome inactivation in development and cancer. FEBS Lett 588:2514–22.

2. Lunyak V V., Rosenfeld MG. 2008. Epigenetic regulation of stem cell fate. Hum Mol Genet 17:R28–36.

3. Sharma S, Kelly TK, Jones PA. 2010. Epigenetics in cancer. Carcinogenesis 31:27–36.

4. Lodish H, Berk A, Matsudaira P, Kaiser CA, Krieger M, Scott MP, Zipursky SL, Darnell J. 2007. Molecular Cell Biology, 6th ed. W.H. Freeman and Company.

5. Funayama R, Ishikawa F. 2007. Cellular senescence and chromatin structure. Chromosoma 116:431–40.

6. Rai TS, Adams PD. 2012. Lessons from senescence: Chromatin maintenance in non-proliferating cells. Biochim Biophys Acta 1819:322–31.

7. Corpet A, Stucki M. 2014. Chromatin maintenance and dynamics in senescence: a spotlight on SAHF formation and the epigenome of senescent cells. Chromosoma 123:423–36.

8. Alper BJ, Lowe BR, Partridge JF. 2012. Centromeric heterochromatin assembly in fission yeast--balancing transcription, RNA interference and chromatin modification. Chromosom Res 20:521–34.

9. Ekwall K. 2007. Epigenetic control of centromere behavior. Annu Rev Genet 41:63–81.

10. Pidoux AL, Allshire RC. 2005. The role of heterochromatin in centromere function. Philos Trans R Soc Lond B Biol Sci 360:569–79.

11. Pidoux AL, Allshire RC. 2004. Kinetochore and heterochromatin domains of the fission yeast centromere. Chromosome Res 12:521–34.

12. Audergon PNCB, Catania S, Kagansky A, Tong P, Shukla M, Pidoux AL, Allshire RC. 2015. Epigenetics. Restricted epigenetic inheritance of H3K9 methylation. Science 348:132–5.

13. Ragunathan K, Jih G, Moazed D. 2015. Epigenetics. Epigenetic inheritance uncoupled from sequence-specific recruitment. Science 348:1258699.

14. Obersriebnig MJ, Pallesen EMH, Sneppen K, Trusina A, Thon G. 2016. Nucleation and spreading of a heterochromatic domain in fission yeast. Nat Commun 7:11518.

15. Muller HJ. 1938. The remaking of chromosomes. Collect Net 8:181–195, 198.

16. McClintock B. 1941. The Stability of Broken Ends of Chromosomes in Zea Mays. Genetics 26:234–82.

17. Watson JD. 1972. Origin of concatemeric T7 DNA. Nat New Biol 239:197–201.

18. Olovnikov AM. 1973. A theory of marginotomy. J Theor Biol 41:181–190.

19. Smogorzewska A, de Lange T. 2004. Regulation of telomerase by telomeric proteins. Annu Rev Biochem 73:177–208.

20. Gottschling DE, Aparicio OM, Billington BL, Zakian VA. 1990. Position effect at S. cerevisiae telomeres: reversible repression of Pol II transcription. Cell 63:751–62.

21. Nimmo ER, Cranston G, Allshire RC. 1994. Telomere-associated chromosome breakage in fission yeast results in variegated expression of adjacent genes. EMBO J 13:3801–11.

22. Castillo AG, Pidoux AL, Catania S, Durand-Dubief M, Choi ES, Hamilton G, Ekwall K, Allshire RC. 2013. Telomeric repeats facilitate CENP-A(Cnp1) incorporation via telomere binding proteins. PLoS One 8:e69673.

23. Kanoh J, Sadaie M, Urano T, Ishikawa F. 2005. Telomere binding protein Taz1 establishes Swi6 heterochromatin independently of RNAi at telomeres. Curr Biol 15:1808–19.

24. Elgin SCR, Grewal SIS. 2003. Heterochromatin: silence is golden. Curr Biol 13:R895–8.

25. Hansen KR, Burns G, Mata J, Volpe TA, Martienssen RA, Bähler J, Thon G. 2005. Global effects on gene expression in fission yeast by silencing and RNA interference machineries. Mol Cell Biol 25:590–601.

26. Hall IM, Shankaranarayana GD, Noma K-I, Ayoub N, Cohen A, Grewal SIS. 2002. Establishment and maintenance of a heterochromatin domain. Science 297:2232–7.

27. Partridge JF, DeBeauchamp JL, Kosinski AM, Ulrich DL, Hadler MJ, Noffsinger VJP. 2007. Functional separation of the requirements for establishment and maintenance of centromeric heterochromatin. Mol Cell 26:593–602.

28. Sadaie M, Iida T, Urano T, Nakayama J-I. 2004. A chromodomain protein, Chp1, is required for the establishment of heterochromatin in fission yeast. EMBO J 23:3825–35.

29. Schalch T, Job G, Noffsinger VJ, Shanker S, Kuscu C, Joshua-Tor L, Partridge JF. 2009. High-affinity binding of Chp1 chromodomain to K9 methylated histone H3 is required to establish centromeric heterochromatin. Mol Cell 34:36–46.

30. Katan-Khaykovich Y, Struhl K. 2005. Heterochromatin formation involves changes in histone modifications over multiple cell generations. EMBO J 24:2138–49.

31. Radman-Livaja M, Ruben G, Weiner A, Friedman N, Kamakaka R, Rando OJ. 2011. Dynamics of Sir3 spreading in budding yeast: secondary recruitment sites and euchromatic localization. EMBO J 30:1012–26.

32. Martins-Taylor K, Dula M Lou, Holmes SG. 2004. Heterochromatin spreading at yeast telomeres occurs in M phase. Genetics 168:65–75.

33. Diede SJ, Gottschling DE. 1999. Telomerase-mediated telomere addition in vivo requires DNA primase and DNA polymerases alpha and delta. Cell 99:723–33.

34. Sunder S, Greeson-Lott NT, Runge KW, Sanders SL. 2012. A new method to efficiently induce a site-specific double-strand break in the fission yeast Schizosaccharomyces pombe. Yeast 29:275–91.

35. Harrison JC, Haber JE. 2006. Surviving the breakup: the DNA damage checkpoint. Annu Rev Genet 40:209–35.

36. Ribeyre C, Shore D. 2012. Anticheckpoint pathways at telomeres in yeast. Nat Struct Mol Biol 19:307–13.

37. Michelson RJ, Rosenstein S, Weinert T. 2005. A telomeric repeat sequence adjacent to a DNA double-stranded break produces an anticheckpoint. Genes Dev 19:2546–59.

38. Lee SS, Bohrson C, Pike AM, Wheelan SJ, Greider CW. 2015. ATM Kinase Is Required for Telomere Elongation in Mouse and Human Cells. Cell Rep 13:1623–1632.

39. Ribeyre C, Shore D. 2013. Regulation of telomere addition at DNA double-strand breaks. Chromosoma 122:159–73.

40. Wang J, Lawry ST, Cohen AL, Jia S. 2014. Chromosome boundary elements and regulation of heterochromatin spreading. Cell Mol Life Sci 71:4841–52.

41. Watson AT, Werler P, Carr AM. 2011. Regulation of gene expression at the fission yeast Schizosaccharomyces pombe urg1 locus. Gene 484:75–85.

42. Grewal SS, Klar AJ. 1997. A recombinationally repressed region between mat2 and mat3 loci shares homology to centromeric repeats and regulates directionality of mating-type switching in fission yeast. Genetics 146:1221–1238.

43. Thon G, Bjerling P, Bunner C, Verhein-Hansen J. 2002. Expression-state boundaries in the mating-type region of fission yeast. Genetics 161:611–622.

44. Wood V, Gwilliam R, Rajandream M-A, Lyne M, Lyne R, Stewart A, Sgouros J, Peat N, Hayles J, Baker S, Basham D, Bowman S, Brooks K, Brown D, Brown S, Chillingworth T, Churcher C, Collins M, Connor R, Cronin A, Davis P, Feltwell T, Fraser A, Gentles S, Goble A, Hamlin N, Harris D, Hidalgo J, Hodgson G, Holroyd S, Hornsby T, Howarth S, Huckle EJ, Hunt S, Jagels K, James K, Jones L, Jones M, Leather S, McDonald S, McLean J, Mooney P, Moule S, Mungall K, Murphy L, Niblett D, Odell C, Oliver K, O’Neil S, Pearson D, Quail MA, Rabbinowitsch E, Rutherford K, Rutter S, Saunders D, Seeger K, Sharp S, Skelton J, Simmonds M, Squares R, Squares S, Stevens K, Taylor K, Taylor RG, Tivey A, Walsh S, Warren T, Whitehead S, Woodward J, Volckaert G, Aert R, Robben J, Grymonprez B, Weltjens I, Vanstreels E, Rieger M, Schäfer M, Müller-Auer S, Gabel C, Fuchs M, Düsterhoft A, Fritzc C, Holzer E, Moestl D, Hilbert H, Borzym K, Langer I, Beck A, Lehrach H, Reinhardt R, Pohl TM, Eger P, Zimmermann W, Wedler H, Wambutt R, Purnelle B, Goffeau A, Cadieu E, Dréano S, Gloux S, Lelaure V, Mottier S, Galibert F, Aves SJ, Xiang Z, Hunt C, Moore K, Hurst SM, Lucas M, Rochet M, Gaillardin C, Tallada VA, Garzon A, Thode G, Daga RR, Cruzado L, Jimenez J, Sánchez M, del Rey F, Benito J, Domínguez A, Revuelta JL, Moreno S, Armstrong J, Forsburg SL, Cerutti L, Lowe T, McCombie WR, Paulsen I, Potashkin J, Shpakovski G V, Ussery D, Barrell BG, Nurse P, Cerrutti L. 2002. The genome sequence of Schizosaccharomyces pombe. Nature 415:871–80.

45. Cromie GA, Rubio CA, Hyppa RW, Smith GR. 2005. A natural meiotic DNA break site in Schizosaccharomyces pombe is a hotspot of gene conversion, highly associated with crossing over. Genetics 169:595–605.

46. Farah JA, Cromie GA, Smith GR. 2009. Ctp1 and Exonuclease 1, alternative nucleases regulated by the MRN complex, are required for efficient meiotic recombination. Proc Natl Acad Sci U S A 106:9356–61.

47. Forsburg SL. 1994. Codon usage table for Schizosaccharomyces pombe. Yeast 10:1045–7.

48. Watson AT, Daigaku Y, Mohebi S, Etheridge TJ, Chahwan C, Murray JM, Carr AM. 2013. Optimisation of the Schizosaccharomyces pombe urg1 expression system. PLoS One 8:e83800.

49. Watt S, Mata J, López-Maury L, Marguerat S, Burns G, Bähler J. 2008. urg1: a uracil-regulatable promoter system for fission yeast with short induction and repression times. PLoS One 3:e1428.

50. Li P, Li J, Li M, Dou K, Zhang M-J, Suo F, Du L-L. 2012. Multiple end joining mechanisms repair a chromosomal DNA break in fission yeast. DNA Repair (Amst) 11:120–30.

51. Wood V, Harris M a., McDowall MD, Rutherford K, Vaughan BW, Staines DM, Aslett M, Lock A, Bähler J, Kersey PJ, Oliver SG. 2012. PomBase: A comprehensive online resource for fission yeast. Nucleic Acids Res 40:695–699.

52. Wang X, Baumann P. 2008. Chromosome fusions following telomere loss are mediated by single-strand annealing. Mol Cell 31:463–73.

53. Webb CJ, Zakian VA. 2008. Identification and characterization of the Schizosaccharomyces pombe TER1 telomerase RNA. Nat Struct Mol Biol 15:34–42.

54. Bairley RCB, Guillaume G, Vega LR, Friedman KL. 2011. A mutation in the catalytic subunit of yeast telomerase alters primer-template alignment while promoting processivity and protein-DNA binding. J Cell Sci 124:4241–52.

55. Murray AW, Claus TE, Szostak JW. 1988. Characterization of two telomeric DNA processing reactions in Saccharomyces cerevisiae. Mol Cell Biol 8:4642–50.

56. Gao Q, Reynolds GE, Wilcox A, Miller D, Cheung P, Artandi SE, Murnane JP. 2008. Telomerase-dependent and -independent chromosome healing in mouse embryonic stem cells. DNA Repair (Amst) 7:1233–49.

57. Wellinger RJ, Zakian VA. 2012. Everything you ever wanted to know about Saccharomyces cerevisiae telomeres: beginning to end. Genetics 191:1073–105.

58. Lydeard JR, Lipkin-Moore Z, Jain S, Eapen V V, Haber JE. 2010. Sgs1 and exo1 redundantly inhibit break-induced replication and de novo telomere addition at broken chromosome ends. PLoS Genet 6:e1000973.

59. Grimm C, Kohli J, Murray J, Maundrell K. 1988. Genetic engineering of Schizosaccharomyces pombe: a system for gene disruption and replacement using the ura4 gene as a selectable marker. Mol Gen Genet 215:81–6.

60. Elgin SCR, Reuter G. 2013. Position-effect variegation, heterochromatin formation, and gene silencing in Drosophila. Cold Spring Harb Perspect Biol 5:a017780.

61. Sabatier L, Ricoul M, Pottier G, Murnane JP. 2005. The loss of a single telomere can result in instability of multiple chromosomes in a human tumor cell line. Mol Cancer Res 3:139–50.

62. Yu S, Graf WD. 2010. Telomere capture as a frequent mechanism for stabilization of the terminal chromosomal deletion associated with inverted duplication. Cytogenet Genome Res 129:265–74.

63. Yatsenko SA, Brundage EK, Roney EK, Cheung SW, Chinault AC, Lupski JR. 2009. Molecular mechanisms for subtelomeric rearrangements associated with the 9q34.3 microdeletion syndrome. Hum Mol Genet 18:1924–36.

64. Osborne EA, Hiraoka Y, Rine J. 2011. Symmetry, asymmetry, and kinetics of silencing establishment in Saccharomyces cerevisiae revealed by single-cell optical assays. Proc Natl Acad Sci U S A 108:1209–16.

65. Hathaway NA, Bell O, Hodges C, Miller EL, Neel DS, Crabtree GR. 2012. Dynamics and memory of heterochromatin in living cells. Cell 149:1447–60.

66. Klar AJS, Ishikawa K, Moore S. 2014. A Unique DNA Recombination Mechanism of the Mating/Cell-type Switching of Fission Yeasts: a Review. Microbiol Spectr 2:1–16.

67. Martienssen R, Moazed D. 2015. RNAi and heterochromatin assembly. Cold Spring Harb Perspect Biol 7:a019323.

68. Allshire RC, Ekwall K. 2015. Epigenetic Regulation of Chromatin States in Schizosaccharomyces pombe. Cold Spring Harb Perspect Biol 7:a018770.

69. Kageyama S, Liu H, Kaneko N, Ooga M, Nagata M, Aoki F. 2007. Alterations in epigenetic modifications during oocyte growth in mice. Reproduction 133:85–94.

70. Ge Z-J, Schatten H, Zhang C-L, Sun Q-Y. 2015. Oocyte ageing and epigenetics. Reproduction 149:R103–14.

71. Moreno S, Klar A, Nurse P. 1991. Molecular genetic analysis of fission yeast Schizosaccharomyces pombe. Methods Enzymol 194:795–823.

72. Forsburg SL. 2011. PombeNet: Drugs.

73. Erler A, Maresca M, Fu J, Stewart AF. 2006. Recombineering reagents for improved inducible expression and selection marker re-use in Schizosaccharomyces pombe. Yeast 23:813–23.

74. Muhlrad D, Hunter R, Parker R. 1992. A rapid method for localized mutagenesis of yeast genes. Yeast 8:79–82.

75. Ray A, Runge KW. 1999. The yeast telomere length counting machinery is sensitive to sequences at the telomere-nontelomere junction. Mol Cell Biol 19:31–45.

76. Hector RE, Ray A, Chen B-R, Shtofman R, Berkner KL, Runge KW. 2012. Mec1p associates with functionally compromised telomeres. Chromosoma 121:277–90.

77. Förstemann K, Höss M, Lingner J. 2000. Telomerase-dependent repeat divergence at the 3’ ends of yeast telomeres. Nucleic Acids Res 28:2690–4.

78. Moser BA, Chang Y, Nakamura TM. 2014. Telomere regulation during the cell cycle in fission yeast. Methods Mol Biol 1170:411–24.

79. Fisher TS, Taggart AKP, Zakian VA. 2004. Cell cycle-dependent regulation of yeast telomerase by Ku. Nat Struct Mol Biol 11:1198–205.

80. Kiely CM, Marguerat S, Garcia JF, Madhani HD, Bähler J, Winston F. 2011. Spt6 is required for heterochromatic silencing in the fission yeast Schizosaccharomyces pombe. Mol Cell Biol 31:4193–204.

81. Kato H, Okazaki K, Iida T, Nakayama J-I, Murakami Y, Urano T. 2013. Spt6 prevents transcription-coupled loss of posttranslationally modified histone H3. Sci Rep 3:2186.

82. Yamada S, Ohta K, Yamada T. 2013. Acetylated Histone H3K9 is associated with meiotic recombination hotspots, and plays a role in recombination redundantly with other factors including the H3K4 methylase Set1 in fission yeast. Nucleic Acids Res 41:3504–17.

83. Oya E, Kato H, Chikashige Y, Tsutsumi C, Hiraoka Y, Murakami Y. 2013. Mediator directs co-transcriptional heterochromatin assembly by RNA interference-dependent and -independent pathways. PLoS Genet 9:e1003677.

84. Kramer KM, Haber JE. 1993. New telomeres in yeast are initiated with a highly selected subset of TG1-3 repeats. Genes Dev 7:2345–56.

